# Medians at the corners, anti-medians at the center: the geometry of breakpoint medians for random signed multichromosomal genomes

**DOI:** 10.64898/2026.10.18.758743

**Authors:** Poly H da Silva, Arash Jamshidpey, David Sankoff

**Affiliations:** Independent Researcher; University of California, Berkeley; University of Ottawa

**Keywords:** Genomes, Breakpoint distance, Medians, Anti-medians

## Abstract

**Purpose:** Breakpoint medians summarize gene order across genomes and are central to ancestral reconstruction. Haghighi and Sankoff (2012) found most breakpoint medians of random genomes near the inputs rather than at balanced positions, while Larlee et al. (2016) observed that balancing adjacency contributions increases the total breakpoint distance, producing anti-medians. We study both phenomena for *k* independent uniform signed multichromosomal genomes on *n* genes.

**Major findings:** Poisson bounds control the median-value deviation from (*k* − 1)*n*, and every median inherits all but *O*_ℙ_(1) of its adjacencies from the inputs. We build candidate genomes by drawing adjacencies from the inputs one at a time. The supply from a fresh input depletes quadratically rather than linearly, making large contributions from multiple inputs harder to combine. We characterize the attainable profiles, the limiting overlaps such a construction can share with each input. At the median scale the constructed genome takes all but *o*_ℙ_(*n*) of its adjacencies from one input, explaining the Haghighi–Sankoff corner-seeking phenomenon. The same characterization quantifies the cost of compromise and provides a game-theoretic explanation of anti-medians. On the Pareto boundary the total overlap is maximal when it all comes from one input and uniquely minimal at equal compromise, where the excess over the median value has a closed form. Numerical experiments illustrate these effects at moderate genome sizes.

**Conclusions:** These results provide a rigorous, quantitative account of breakpoint median and anti-median geometry in random signed genomes, showing why attainable median-scale constructions favor a single input and what balanced compromise costs.

## 1 Introduction

Gene order carries evolutionary information that complements sequence data, since rearrangements such as inversions, translocations, fusions and fissions reshuffle genes along and between chromosomes, and the resulting differences in gene order can be used to study evolutionary relationships. One of the simplest and most widely studied measures of gene-order dissimilarity is the breakpoint distance [13], which compares the adjacencies present in two genomes. For three or more genomes, a natural representative is a breakpoint median, a genome minimizing the sum of its breakpoint distances to the inputs. This optimization principle motivates the use of medians in ancestral genome reconstruction and gene-order phylogenetics.

The breakpoint median problem was introduced by [14], who related it to the Traveling Salesman Problem (TSP), and it is NP-hard for unichromosomal unsigned genomes [2]. More generally, median problems have been studied under various genomic distances, including breakpoint, reversal, DCJ and SCJ distances, with computational properties depending strongly on both the distance and the genome model [2, 3, 15, 8, 7, 16, 18]. Beyond computational complexity, however, there is a more basic geometric question. What does a typical median look like, and how is it positioned relative to the genomes it summarizes?

Breakpoint medians are often interpreted as ancestral genomes, and one therefore expects a median to combine information from several inputs rather than nearly reproduce one of them. Simulations by [9] challenged this expectation. For *k* random unichromosomal genomes on *n* genes, they observed a median value close to (*k* −1)*n*, essentially the total distance obtained by choosing one of the inputs itself. Moreover, most heuristic medians lay very close to one input genome, near a “corner” of genome space. Only a small proportion appeared away from the corners, some roughly equidistant from all inputs. Such intermediate medians are particularly relevant to ancestral reconstruction, since they could represent genomic information from several lineages in a balanced way. This corner-seeking behavior is known as the Haghighi–Sankoff conjecture.

One reason why finding medians far from the corners is difficult is that genome space endowed with breakpoint distance is non-geodesic. Consequently, noncorner medians cannot in general be reached by moving gradually along shortest paths from one input toward the others [10]. [10] partially proved the Haghighi–Sankoff conjecture by establishing the asymptotic median value (*k* −1)*n* and introduced partial geodesics as a means of reaching non-trivial medians far from the corners, termed “accessible genomes.” They also showed that, for random genomes, a median contains only a small number of adjacencies absent from all inputs.

A natural route to noncorner medians is therefore to draw adjacencies from several inputs at once. [12, 11] tested this idea on random signed genomes, seeking a genome that inherits the same proportion *x* of adjacencies from each of the *k* inputs, with *kx* as close to 1 as possible. The outcome was the opposite of what was expected. The more evenly the adjacencies were shared, the smaller the achievable *x*, and the farther the resulting genome was from the inputs in total. They called such genomes, equidistant from the inputs but far from optimal, anti-medians.

Sampling adjacencies from different inputs, however, gives little control over whether the chosen adjacencies can coexist in a single genome. To address this, [4] classified the adjacencies of a random genome relative to a previously selected set, making it possible to count how many remain usable. In related work, [5] introduced a branching algorithm that identifies breakpoint median genomes and used it to enumerate, for the first time, all exact medians of random unichromosomal unsigned genomes. [6] develop this into a family of tree-based algorithms whose leaves contain all exact and approximate medians in a search space far smaller than the space of all genomes, together with sampling procedures that target balanced near-medians. They show that the number of medians grows exponentially in the number of genes, and theirs are the first methods able to produce medians away from the corners, in particular medians equidistant from the inputs. Although presented for unsigned unichromosomal genomes, the algorithms apply equally to multichromosomal genomes. For unsigned unichromosomal genomes, we expect the framework of [4] to play a key role in settling the conjecture. Signed and unsigned genomes, however, obey substantially different compatibility constraints, and these differences make the signed multichromosomal model considered here more tractable.

In this paper we give a rigorous account of both phenomena, the corner-seeking behavior of medians and the emergence of anti-medians, for *k* independently and uniformly sampled signed multichromosomal genomes. For simplicity, all chromosomes are taken to be circular, and the methods extend naturally to linear chromosomes through a standard telomere-capping construction. We first show that the deviation of the median value from (*k* −1)*n* remains stochastically bounded, with fluctuations controlled by explicit Poisson bounds, and that every median contains at most a stochastically bounded number of adjacencies absent from all inputs.

To find medians, we build candidate medians sequentially, drawing adjacencies from the input genomes one genome at a time as in [12, 11]. At each stage only adjacencies whose two extremities are still free can be added, so that the selection remains part of a single genome. The key observation is that the supply of usable adjacencies depletes quadratically. Once a proportion *s* of a candidate genome has been fixed, a fresh random input contains asymptotically only a proportion (1 −*s*)^2^ of usable adjacencies, while a proportion 1 −*s* ≥ (1 −*s*)^2^ of the genome remains to be filled. Combining substantial contributions from several inputs therefore becomes progressively harder. The corresponding depletion is weaker in the unsigned setting [4], and it is the stronger depletion in the signed model that makes a rigorous treatment possible here.

We formalize this construction as an adapted sequential sampling scheme, in which the choices made at each stage may depend only on the input genomes already revealed. The selected adjacencies are then completed to a genome by pairing the remaining extremities uniformly at random. We characterize exactly which limiting overlap profiles, that is, which asymptotic proportions of adjacencies shared with each of the *k* inputs, are attainable. In particular, if these proportions sum to one, as required asymptotically for a median, the only attainable profiles are the corners of the probability simplex. Thus, every approximate median obtained in this way draws almost all of its adjacencies from a single input genome, which may depend on the realization, and approaches that genome in normalized breakpoint distance. This yields a rigorous proof of the Haghighi–Sankoff corner-seeking phenomenon for signed multichromosomal genomes through attainable overlap profiles.

The same characterization also quantifies the cost of compromise and gives a natural game-theoretic interpretation of the anti-medians observed in [11]. We regard the *k* input genomes as players whose payoffs are the proportions of their adjacencies retained in the constructed genome. The total payoff is asymptotically *k* minus the normalized total breakpoint distance to the inputs, so maximizing it amounts to approaching the median value. For general *k*, we characterize the Pareto-optimal boundary of the feasible payoff region and show that the total payoff is maximized precisely at the corners and uniquely minimized at the equal-compromise point (*a*_*k*_, …, *a*_*k*_), for which we obtain a closed-form expression. For three genomes, we further determine the complete trade-off between a prescribed minimum overlap *q* with each input and the largest attainable total overlap. Equivalently, the normalized asymptotic excess over the median value increases continuously from 0 at *q* = 0 to 1/4 at the equal-compromise point *q* = 1/4. Thus, among Pareto-optimal configurations, full compromise incurs the largest distance penalty, providing a quantitative explanation for the anti-median behavior observed in simulations.

Finally, we examine how these asymptotic results are reflected at finite genome sizes. Simulations for *n* = 100, …, 800 closely follow the theoretical compromise costs, both for three genomes and for equal compromise with *k* = 2, …, 6. Sequential sampling schemes also reproduce the attainable-profile geometry, with high success probabilities typically occurring inside the limiting region and low probabilities outside it. Thus, both the cost of compromise and the attainable-profile geometry are already visible at moderate genome sizes.

The remainder of the paper is organized as follows. Section 2 introduces the genome model and establishes the probabilistic results for random break-point medians. Section 3 develops the adapted sequential sampling framework, characterizes the attainable overlap region, and proves the corner-seeking result under sequential sampling. Section 4 treats the input genomes as cooperating players, characterizes the Pareto-optimal boundary of their feasible payoffs, and shows that the total payoff on this boundary is maximal at the corners and uniquely minimal at equal compromise, the anti-median, for which the excess over the median value has a closed form. Section 5 presents the finite-size numerical experiments, and Section 6 concludes with a discussion.

## 2 Asymptotic Structure of Breakpoint Medians

Although the methods in this paper extend readily to multichromosomal genomes with linear and circular chromosomes, using standard capping arguments, for simplicity we assume that all chromosomes are circular. With no telomeres, a signed multichromosomal genome with circular chromosomes can be represented as a perfect matching on the set of vertices ±[*n*] := {±1, ±2, …, ±*n*}. For genes labeled 1, …, *n*, we denote the two extremities of gene *i* by −*i* and +*i*, and the signs may be interpreted as encoding gene polarity (strandedness). Each edge of the matching represents a gene adjacency. Allowing arbitrary numbers and sizes of chromosomes places no restrictions on the associated perfect matching. For instance, an edge { −*i*, +*i*} means that the genome contains a circular chromosome of size 1, with gene content *i*. We denote by G_*n*_ the set of all perfect matchings on ± [*n*]. More generally, the gene content of a genome *G* may be any finite subset ±*V* (*G*) ⊆ [*n*]. Its extremities are ± *V* (*G*) := *V* (*G*) ∪ (−*V* (*G*)), and we represent *G* as the perfect matching *G* = (±*V* (*G*), *E*(*G*)). In particular, for *G* ∈ G_*n*_, ±*V* (*G*) = ± [*n*].

Breakpoint distance (bp-distance), introduced in [13], is often used to measure the dissimilarity between two genomes. More specifically, for a pair of genomes *G, G*′, the breakpoint distance *d*(*G, G*′) can be defined by *d*(*G, G*′) = |*E*(*G*) Δ *E*(*G*′)|/2, where Δ denotes the symmetric difference between sets and | · | denotes the cardinality of the set. In particular, for *G, G*′ ∈ G_*n*_, this formula reduces to *d*(*G, G*′) = *n* − |*E*(*G*) ∩ *E*(*G*′)|.

Letting G be the space of all perfect matchings on ±*S* := *S* ∪(−*S*), for finite subsets *S* ⊂ ℒ, (G, *d*) is a metric space. In addition, for any *n* ∈ ℒ, (G_*n*_, *d*) is also a metric space. Let **S**_±*n*_ be the group of permutations on ±[*n*] with the function composition as the group action. We can define the action of **S**_±*n*_ on G_*n*_ by *σ* · *G* ∈ G_*n*_ for *σ* ∈ **S**_±*n*_ and *G* ∈ G_*n*_, where ±*V* (*σ* · *G*) = ±[*n*], and *E*(*σ* · *G*) = {{*σ*(*i*), *σ*(*j*)} : {*i, j*} ∈ *E*(*G*)}. That is, the action relabels the gene extremities via the permutation *σ* ∈ **S**_±*n*_.

Indeed, the action of **S**_±*n*_ on G_*n*_ defined above is distance preserving, i.e., for any *G, G*′ ∈ G_*n*_ and *σ* ∈ **S**_±*n*_, *d*(*σ* · *G, σ* · *G*′) = *d*(*G, G*′). We say that the bp-distance is left-invariant under the action of **S**_±*n*_ on G_*n*_.

We begin by analyzing the distribution of common adjacencies between independent random genomes. This provides a probabilistic baseline for genome comparison and plays a key role in understanding the structure and typical behavior of breakpoint medians in later sections.

Let *G* and *G*′ be two independent genomes chosen uniformly at random from G_*n*_. Let *o*_*n*_ ∈ G_*n*_ be the perfect matching on ±[*n*] defined by *E*(*o*_*n*_) = {{−*i*, +*i*} : *i* ∈ [*n*]}. Let (*G*′)^−1^ ⊂ **S**_±*n*_ denote the set of all permutations *σ* such that *σ* · *G*′ = *o*_*n*_, and, conditional on *G*′, let 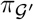 be chosen uniformly from (*G*′)^−1^, independently of *G*. Since, for every fixed *σ* ∈ **S**_±*n*_, the action *G* ↦→ *σ* ·*G* is a bijection of G_*n*_, the uniform distribution on G_*n*_ is invariant under this action. Hence 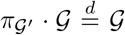. Moreover, by left-invariance of the breakpoint distance, 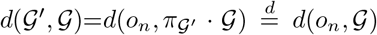, where 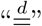 indicates equivalence in distribution. This reduction allows us to replace a two-genome comparison with a one-genome problem relative to a fixed reference, simplifying the analysis. More precisely, to study the bp-distance of two independent random genomes, it suffices to study the bp-distance of a uniformly random genome *G* ∈ G_*n*_ and the reference genome *o*_*n*_ ∈ *G*_*n*_. Note that the choice of *o*_*n*_ is arbitrary and any other fixed perfect matching in G_*n*_ can serve as the reference genome. To simplify notation, for the set of genomes *V* = {*G*_1_, …, *G*_*m*_}, let 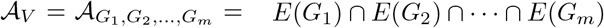.

It is well known that for two independent uniformly sampled genomes 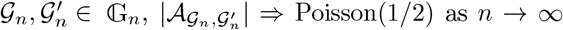 see, for example, [17]. We use this pairwise Poisson limit below to analyze the medians of *k* independent uniformly sampled genomes.

The geometric medians were used in comparative genomics for the first time by [14]. They have extensive applications in gene-order phylogenetics. A breakpoint (bp) median of a set of genomes *A* = {*G*_1_, *G*_2_, …, *G*_*k*_} ⊆ G_*n*_ is a genome *G* ∈ G_*n*_ that minimizes the total breakpoint distance function 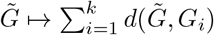. The minimum value of the function is called the bp-median value of *A*. We denote by *M*_*n*_(*A*) = *M* (*A*) = *M* (*G*_1_, …, *G*_*k*_), the set of all bpmedians of *A*, and by *µ*_*n*_(*A*) = *µ*(*A*) = *µ*(*G*_1_, …, *G*_*k*_), the median value of *A*. The median of a single genome *G* is itself. For two genomes *G*_1_, *G*_2_ ∈ G_*n*_, the triangle inequality gives *µ*(*G*_1_, *G*_2_) = *d*(*G*_1_, *G*_2_), and *G* ∈ *M* (*G*_1_, *G*_2_) if and only if 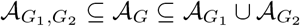 (see also [10] for the unsigned case). In addition, for *k* genomes *A* = {*G*_1_, *G*_2_, …, *G*_*k*_} ⊆ G_*n*_, for any fixed *i* ∈ [*k*],

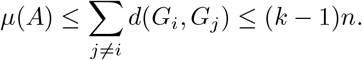

As mentioned above, for *k* perfect matchings *G*_1_, *G*_2_, …, *G*_*k*_, chosen independently and uniformly at random from G_*n*_, the pairwise distances satisfy *d*(*G*_*i*_, *G*_*j*_) = *n* −*O*_ℙ_(1), with Poisson(1/2)-type fluctuations in the number of common adjacencies. Consequently, studying the medians and median value of *G*_1_, …, *G*_*k*_ is similar to studying those of deterministic genomes *G*_1_, …, *G*_*k*_ with pairwise maximum distance *d*(*G*_*i*_, *G*_*j*_) = *n* for *i* ≠ *j*. Indeed, if *A* = {*G*_1_, …, *G*_*k*_} ⊂ G_*n*_ satisfies *d*(*G*_*i*_, *G*_*j*_) = *n* for all *i* ≠ *j*, then the adjacency sets of the inputs are pairwise disjoint and we have *µ*(*A*) = (*k* − 1)*n*, and 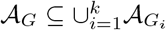 for every *G* ∈ *M* (*A*). Indeed, 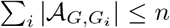, with equality at each input genome, and equality for a median forces all its adjacencies to lie in the union. We now make this comparison precise.

To this end, for *A* = {*G*_1_, …, *G*_*k*_} ⊆ G_*n*_, let 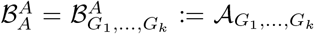. Then, for any *j* = 1, · · ·, *k*, define

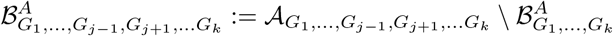

Continuing inductively, for any 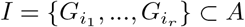, we set

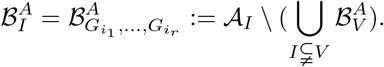

In other words, 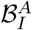 consists of all adjacencies that are present in every genome in *I*, but absent from every genome in *A I*. The sets 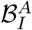 decompose shared adjacencies according to the subsets of genomes in which they appear.

Let *U*_*n,k*_, *L*_*n,k*_ : (G_*n*_)^*k*^ → ℤ_+_ be defined by

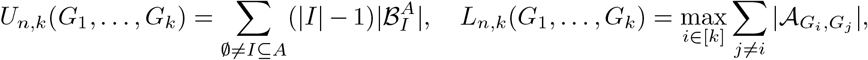

where in the latter, for each fixed *i* ∈ [*k*], the inner sum is taken over all *j* ∈ [*k*] with *j* ≠ *i*. The outer maximum then selects the largest value of this sum over all choices of *i*.

The next result provides explicit upper and lower bounds on how far the median value can fall below the maximal value (*k* −1)*n*. These bounds depend only on how adjacencies are shared among subsets of the input genomes. The proof is given in Appendix A.

### Theorem 1.

*For any k* ≥ 3, *and any set A* = {*G*_1_, …, *G*_*k*_} *of k distinct perfect matchings in* G_*n*_, *we have*

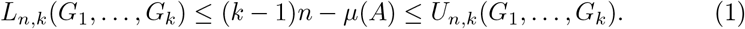

In other words, Theorem 1 shows that the deviation of the median value from its maximum is governed by the shared adjacencies among the input genomes. For random genomes, such shared structures occur only sparsely, leading to tight probabilistic control over the median value.

We now apply this on a set of *k* independent random perfect matchings *A* = {*G*_1_, …, *G*_*k*_} ⊂ G_*n*_. As mentioned, this is similar to the case of *k* non-random perfect matchings with pairwise distance *n*. Let *α*^*^ and *β*^*^ be two independent Poisson random variables,

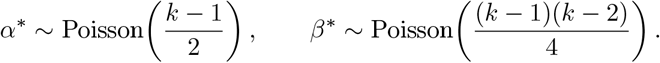

The following lemma collects the common-adjacency asymptotics that will be used later. Its proof is deferred to Appendix A. Here and below, ⇒ denotes convergence in distribution.

### Lemma 1.

*Let k* ≥ 3, *and let A* = {*G*_1_, …, *G*_*k*_} ⊆ G_*n*_ *be a set of genomes chosen uniformly and independently at random from* G_*n*_. *Then, as n* → ∞, |*A*_*I*_ | ⇒ 0 *for every I* ⊆ *A with* |*I*| ≥ 3, *and for any i* ∈ [*k*],

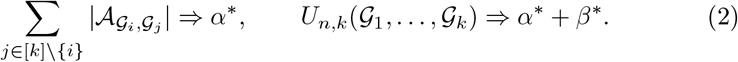

We say a nonnegative integer-valued random variable *Z* is stochastically dominated by *Y, Z* ⪯_*st*_ *Y*, if for every *r* ∈ ℤ_+_, ℙ(*Z > r*) ≤ ℙ(*Y > r*).

The following theorem provides stochastic bounds on the deviation of the median value of *k* independent random genomes from (*k* −1)*n*. The proof is given in Appendix A.

### Theorem 2.

*For a given k* ≥ 3, *let* 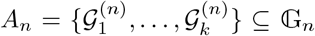 *be k perfect matchings chosen independently and uniformly at random from* G_*n*_. *Then for every r* ∈ ℤ_+_,

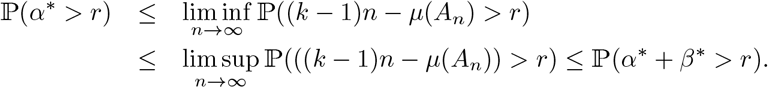

*Furthermore*, 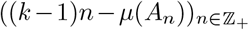 *is tight and hence every subsequence has a further subsequence that converges in distribution. For every subsequential limit X*^*^, *we have α*^*^ ⪯_*st*_ *X*^*^ ⪯_*st*_ *α*^*^ + *β*^*^.

The previous theorem shows that the deviation (*k* −1)*n* −*µ*(*A*_*n*_) remains stochastically bounded as *n* → ∞. In particular, the median value differs from its maximal possible value (*k* −1)*n* only by a random quantity of order *O*_ℙ_(1). The next result shows that this deviation becomes negligible under any diverging normalization. The proof is deferred to Appendix A.

### Theorem 3.

*Under the assumptions of Theorem 2, for any diverging sequence* (*a*_*n*_)_*n*≥1_ *of strictly positive numbers, a*_*n*_ → ∞, *as n* → ∞, *we have*

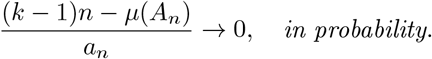

While the previous result concerns the median value, it does not yet describe the adjacency structure of the median genome itself. The following theorem addresses this question by showing that, with high probability, a median genome introduces only a vanishing proportion of adjacencies not present in the input genomes. The proof is given in Appendix A.

For *k, n* ≥ 1 and *G*_1_, …, *G*_*k*_ ∈ G_*n*_, define

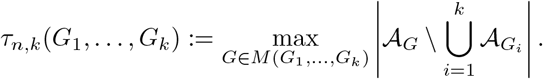

### Theorem 4.

*For a given k* ≥ 1, *let* 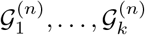 *be k perfect matchings chosen independently and uniformly at random from* G_*n*_. *For every diverging sequence* (*a*_*n*_)_*n*≥1_ *of strictly positive numbers*,

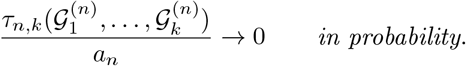

These results show that the bp-median value is asymptotically close to (*k* −1)*n*, while any median introduces only a vanishing proportion of new adjacencies.

## 3 Medians Tend to Corners

In this section, we study the positions of medians of *k* independent random perfect matchings (genomes) in G_*n*_, where *k* ≥ 2 is fixed. As shown in the previous section, the median value is (*k* −1)*n* −*O*_ℙ_(1), and the number of adjacencies of a median that do not belong to the union of the input genomes becomes negligible. More precisely, after division by any sequence tending to infinity, this quantity converges to zero in probability.

While the previous section showed that the median value of random genomes is close to its maximum possible value, this does not by itself determine where a median genome is located in the genome space. Let 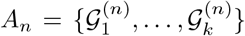 be a collection of genomes, sampled uniformly and independently at random from G_*n*_. Given nonnegative weights *x*_1_, …, *x*_*k*_ with *x*_1_ + · · ·+ *x*_*k*_ = 1, we ask whether one can construct a genome whose normalized overlaps with the input genomes approach *x*_1_, …, *x*_*k*_, and in particular whether a median-scale construction can combine substantial proportions of adjacencies from multiple different inputs. We denote by Δ_*k*_ the (*k* − 1)-dimensional probability simplex, Δ_*k*_ := {(*x*_1_, …, *x*_*k*_) ∈ ℝ^*k*^ : *x*_*i*_ ≥ 0 for all *i, x*_1_ + · · · + *x*_*k*_ = 1}. Thus, Δ_*k*_ parametrizes the limiting normalized overlap profiles of the potential approximate medians. Points in the interior of Δ_*k*_ represent genuinely mixed profiles, in which substantial proportions of adjacencies are inherited from multiple input genomes, whereas the vertices *e*_1_, …, *e*_*k*_ represent profiles whose adjacency content is asymptotically concentrated in a single input genome.

The probabilistic mechanism has a simple geometric interpretation. Suppose that a proportion *s* of the adjacencies of a candidate genome has already been fixed. A fresh independent input genome then contains asymptotically only a proportion (1 −*s*)^2^ of adjacencies whose two extremities remain available. Thus the usable fraction decreases quadratically, rather than linearly, as contributions are accumulated. Distributing a substantial fraction of a candidate genome among several independent inputs therefore progressively reduces the supply of adjacencies available from subsequent inputs. This depletion of compatible choices creates a stochastic pressure toward the corners of Δ_*k*_. Constructions concentrated around one input avoid successive quadratic losses, while mixed constructions become increasingly constrained.

The compatibility mechanism also has a combinatorial interpretation. Fix the number of adjacencies to be taken from each input genome, and call a choice of these adjacencies a compatible selection if the chosen blocks are pairwise vertex-disjoint, so that together they form part of a single genome. Since a uniformly chosen block of a uniform random matching is itself a uniform matching of that size on ±[*n*], averaging over the inputs leaves a pure counting problem, and compatible selections can be counted either one input at a time, multiplying the matchings still available at each stage, or all at once. The two counts agree, so the number of compatible selections does not depend on the order in which the inputs are taken. Correspondingly, after averaging over the inputs, the sequential procedure with uniform choices at each stage, conditioned on succeeding at every stage, produces a uniformly random compatible selection, and so does a quite different procedure, namely choosing the prescribed number of adjacencies from each input independently, without imposing compatibility, and keeping the outcome only when the chosen blocks happen to be pairwise vertex-disjoint. The equivalence is proved in Appendix A.7. The order of exposure can change how often the sequential procedure succeeds, but not the law of the selection it produces given success. For each fixed order, the success probability of the scheme depends only on the prescribed numbers and *n*. Sequential sampling is therefore a way of counting compatible selections, not only a way of producing one.

Counting also shows that balanced selections are less likely to be compatible. For a fixed total number of selected adjacencies, balancing the prescribed contributions decreases the proportion of compatible selections among all choices of those sizes, even though the absolute number of compatible selections increases. Thus, this proportion is largest when the selection is concentrated on a single input, near corners, and smallest when it is distributed as evenly as possible. The lower compatibility of balanced selections is therefore a property of the configurations themselves, not of the order of exposure or of any particular selection rule. Below, we use the quadratic free-edge law to characterize attainable profiles and establish the corresponding median-scale corner geometry.

We say that a sequence of random variables (*Z*_*n*_)_*n*≥0_ converges to a random variable *Z* in *ℙ*, denoted 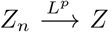, if E(|*Z* − *Z*|^*p*^) → 0, *n* → ∞. It is well known that convergence in *L*^2^ implies convergence in probability, and the latter implies convergence in distribution. In what follows, we use this fact through a precise control of certain edge counts in random genomes. A key ingredient is understanding how many edges of a random genome can avoid a prescribed set of gene extremities.

### Proposition 1.

*Let m* = *m*_*n*_ {0, …, *n*} *be such that m/n* → *x* ∈ [0, 1] *as n* → ∞, *and let R*_*n*_ ⊆ ±[*n*] *be any given set with* |*R*_*n*_| = 2*m. Let G*^(*n*)^ *be a perfect matching selected uniformly at random from* G_*n*_, *and let F*_*n*_ = *F*_*n*_(*R*_*n*_) *be the number of edges in E*(*G*^(*n*)^) *whose two vertices both belong to* ±[*n*] \ *R*_*n*_. *Then*

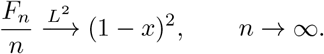

Proposition 1 shows that, asymptotically, the number of edges that remain available after fixing a positive fraction of the gene extremities concentrates around a deterministic value. In particular, once a proportion *x* of the adjacencies has been fixed, the proportion of adjacencies of a new independent genome whose two extremities remain free approaches (1 − *x*)^2^.

### Remark 1.

*As the distribution of F*_*n*_ *does not depend on the specific choice of R*_*n*_ *but only on its cardinality, we may regard F*_*n*_ *as a function of m, writing F*_*n*_ = *F*_*n*_(*m*). *The same conclusion holds conditionally when R*_*n*_ *is random but independent of G*^(*n*)^, *provided its cardinality is prescribed as above. We will also use the uniform first-order consequence*

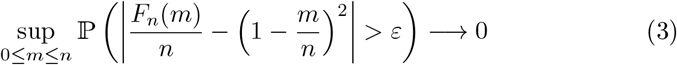

*for every ε >* 0, *as n*→ ∞. *Indeed, if this failed, one could choose a sequence of sizes for which the deviation probability remains bounded away from zero. A subsequence of the corresponding normalized sizes would converge in* [0, 1], *contradicting Proposition 1*.

For a genome *G* ∈ G_*n*_ and a set *R* ⊆ ±[*n*], we say that an edge of *G* is *free with respect to R* if both of its vertices belong to ±[*n*] \ *R*. Thus, Proposition 1 describes the asymptotic number of free edges of an independent random genome relative to a prescribed set of occupied extremities.

We now describe a sequential procedure for sampling gene adjacencies from the input genomes. The inputs are considered one at a time in a fixed order, and at each stage adjacencies of the current input are added to a growing partial genome.

### Definition 1

(Adapted sequential sampling scheme). *Fix an order σ*(1), …, *σ*(*k*) *of the input genomes and start from* 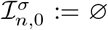. *At stage r* = 1, …, *k, select a set I*_*n,σ*(*r*)_ *of adjacencies of 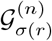 that are free with respect to the extremities of* 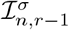, *and set* 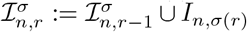. *The selection at stage r may depend on the input genomes already considered and on all earlier selections, and may use randomness independent of the input genomes, but it may not depend on any input genome not yet considered*.

Since only free adjacencies are added, each 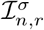 is a matching and the sets *I*_*n*,1_, …, *I*_*n,k*_ are pairwise vertex-disjoint. Their sizes may be random, and some of them may be empty. A formal probabilistic formulation of this definition is given in Appendix A.8. Here, “adapted” means that the choices made at each stage cannot depend on input genomes that have not yet been considered. For brevity, we henceforth refer to such a procedure as a *sequential sampling scheme*.

An important special case prescribes the sizes in advance. Fix target counts *m*_*n*,1_, …, *m*_*n,k*_ ∈ {0, …, *n*} with 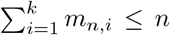, independently of the input genomes. A *prescribed-count scheme* selects exactly *m*_*n,σ*(*r*)_ free adjacencies at stage *r*. If fewer are available, it fails and selects nothing at that or any later stage. It succeeds if all *k* stages are completed. We write 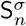 for this success event and 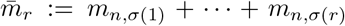. After *r* successful stages exactly 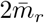 extremities are occupied, and the next input is independent of everything selected so far. Hence, with *F*_*n*_(*m*) as in the remark following Proposition 1,

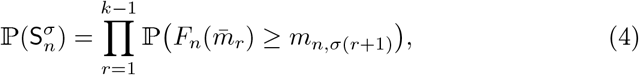

whatever rule is used to choose the adjacencies at each stage.

The adaptedness restriction allows Proposition 1 to be applied at each stage. After the first *r* stages exactly 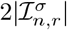 extremities are occupied. Given the earlier inputs and choices, the next input remains an independent uniformly distributed genome. Thus, if the selected proportion approaches *s*, the proportion of free adjacencies supplied by the next input approaches (1 −*s*)^2^. Consequently, if *m*_*n,i*_*/n* → *x*_*i*_ for every *i*, a prescribed-count scheme can succeed with probability tending to one only if

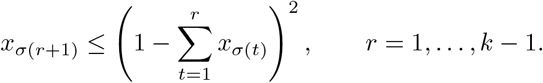

These inequalities express the limiting compatibility constraints. We next define attainability and show that these constraints characterize it exactly.

### Definition 2.

*Let x* = (*x*_1_, …, *x*_*k*_) ∈ [0, 1]^*k*^. *We say that x is σ*-attainable *if there exist deterministic target counts m*_*n*,1_, …, *m*_*n,k*_ *satisfying m*_*n,i*_*/n* → *x*_*i*_, *as n* → ∞, *for every i* ∈ [*k*], *and prescribed-count schemes in the order σ whose probability of succeeding through all k stages tends to one as n* → ∞ . *We call x* attainable *if it is σ-attainable for at least one permutation σ of* [*k*].

For a fixed order *σ*, write 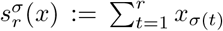, for *r* = 1, …, *k*, and set 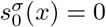. Define

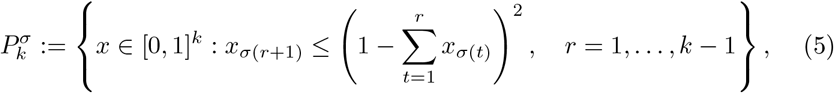

and

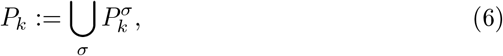

where the union is over all permutations *σ* of [*k*]. For *x* ∈ [0, 1]^*k*^ we write *u*(*x*) := *x*_1_ + · · · + *x*_*k*_ for the *total overlap*, so that 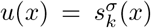 for every permutation *σ* of [*k*].

### Remark 2.

*Every x* ∈ *P*^*σ*^ *satisfies u*(*x*) ≤ 1. *Indeed, writing* 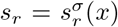, *we have s*_1_ ≤ 1, *and whenever* 0 ≤ *s*_*r*_ ≤ 1 *the constraint gives x*_*σ*(*r*+1)_ ≤ (1 − *s*_*r*_)^2^ ≤ 1 − *s*_*r*_, *so s*_*r*+1_ ≤ 1, *and induction over r* = 1, …, *k* − 1 *gives u*(*x*) = *s*_*k*_ ≤ 1.

The following result shows that the quadratic constraints exactly characterize the limiting profiles attainable by sequential sampling. For *y* ∈ ℝ^*k*^ and *B* ⊆ ℝ^*k*^, let dist(*y, B*) := inf_*z*∈*B*_ max_*i*_ |*y*_*i*_ − *z*_*i*_|.

### Theorem 5.

*For every permutation σ of* [*k*], *a profile x* ∈ [0, 1]^*k*^ *is σ-attainable if and only if x* 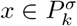. *Consequently, the set of attainable profiles is exactly P*_*k*_. *Moreover, for every sequence of sequential sampling schemes* 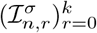, *n* ≥ 1, *in the order σ, as n* → ∞,

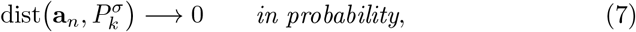

*where* **a**_*n*_ := *n*^−1^(|*I*_*n*,1_|, …, |*I*_*n,k*_|) *is the selected profile, with* 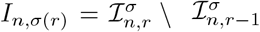.

Both parts rest on the fact that, at each stage, the number of selected adjacencies cannot exceed the number of free adjacencies supplied by the current input, which Proposition 1 controls given the earlier stages. This yields the second part, and necessity follows by applying it to prescribed-count schemes. Conversely, if all the quadratic inequalities above hold strictly, Proposition 1 implies that each stage succeeds with probability tending to one, and therefore so does the full sequential construction. Boundary profiles are obtained by choosing target proportions inside the attainable region and letting them approach the desired boundary profile sufficiently slowly. The full proof is given in Appendix A.8. The second part shows that the quadratic constraints bind every sequential sampling scheme, not only those with counts fixed in advance. Counts that are random, or that depend on the inputs already considered, still yield a selected profile whose distance to 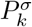 tends to zero in probability.

Theorem 5 does more than record necessary constraints. It shows that every profile in *P*_*k*_ is attainable, including every profile on the boundary. In particular, a profile for which one of the quadratic constraints holds with equality is still attainable. At such a profile the supply of free adjacencies matches the demand exactly in the limit, but at each *n* the target counts can be chosen strictly inside the region and allowed to approach the boundary profile slowly enough that every stage still succeeds with probability tending to one. The region therefore contains genuinely mixed profiles with substantial contributions from several inputs, and these are realized with probability tending to one rather than only permitted by the constraints. Its Pareto-optimal boundary describes the maximal compatible trade-offs between these contributions and plays a central role in the compromise analysis of the next section.

The number of adjacencies selected from an input and the number of adjacencies the final genome shares with that input need not agree. The second can be larger, because an adjacency selected from one input may also belong to another, and because an edge added to complete the partial matching may belong to one of the inputs. We complete the selection uniformly. Conditionally on the input genomes and all choices of the scheme, the 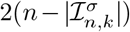 unmatched extremities are paired by a uniformly random perfect matching. We call the resulting genome 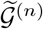 the *uniform completion* of the scheme.

### Proposition 2

(Realization). *Let* 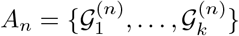 *consist of k* ≥ 2 *independent uniformly distributed random genomes in* G_*n*_, *and let* 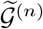 *be the uniform completion of a sequence of sequential sampling schemes* 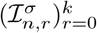, *n* ≥ 1, *in some order σ, with increments* 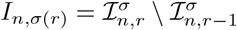, *r* = 1, …, *k. Then*

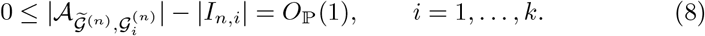

*In particular, for every x* ∈ *P*_*k*_ *there is such a sequence of schemes for which* 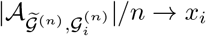 *in probability for every i, as n* → ∞.

The proof is given in Appendix A. The case of boundary profiles *x* is included. Since *d*(*G, H*) = *n* − |*A*_*G,H*_ | and 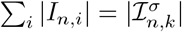, summing (8) over *I* yields

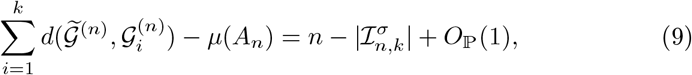

where we used (*k* − 1)*n* − *µ*(*A*_*n*_) = *O*_ℙ_(1). This holds by Theorem 2 for *k* ≥ 3, and for *k* = 2 because 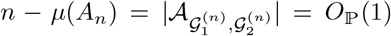. The excess over the median value is therefore, up to a stochastically bounded error, the number of edges added by the completion. A completed genome with limiting profile *x* ∈ *P*_*k*_ has excess (1 − *u*(*x*))*n* + *o*_ℙ_(*n*), so it is an approximate median in the sense of Theorem 6 below if and only if *x* ∈ Δ_*k*_.

For example, the balanced profile (1/4, 1/4, 1/4) belongs to *P*_3_. The corresponding completed genome shares asymptotically a quarter of its adjacencies with each of three independent inputs, and its excess over the median value is *n/*4 + *o*_ℙ_(*n*). Thus, under sequential sampling, mixed intermediate genomes are attainable, but at a macroscopic cost. The Pareto boundary analysis in the next section makes this trade-off quantitative for general *k* and for intermediate levels of compromise. At the median scale no compromise survives under sequential sampling, as the following theorem shows.

### Theorem 6.

*We have P*_*k*_∩Δ_*k*_ = {*e*_1_, …, *e*_*k*_}. *Moreover, let* 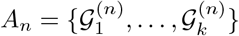 *consist of k* ≥ 2 *independent uniformly distributed random genomes in* G_*n*_, *and let* 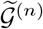 *be the uniform completion of a sequence of sequential sampling schemes* 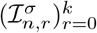, *n* ≥ 1, *in some order σ. If* 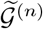 *is an approximate median, that is*,

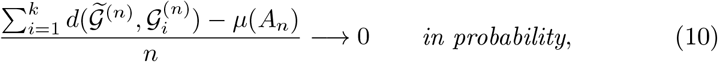

*or in particular if P* 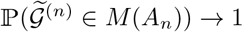 then, in probability, as n → ∞,

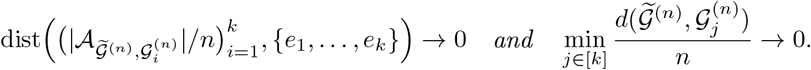

The proof is given in Appendix A. The geometric identity follows from the quadratic constraints. When the coordinates sum to one, the constraint on the last coordinate in the sampling order forces it to be zero or one, and repeating the argument for the preceding coordinates leaves exactly one coordinate equal to one. For the second part, (9) shows that an approximate median leaves only *o*_ℙ_(*n*) edges to the completion, and the second part of Theorem 5 then forces the selected profile, and with it the overlap profile, toward a corner. The theorem does not assert that the overlap profile converges. Only its distance to the set of corners tends to zero, so the corner it approaches, and hence the input, may depend on the realization. The uniformity of the completion is used only to obtain (9), by ensuring that the completion edges add only *O*_ℙ_(1) overlap with the inputs. Some restriction of this kind is necessary. Without it, a scheme that selects nothing could be completed to an arbitrary median of *A*_*n*_, and the theorem would then assert that every median lies close to one of the inputs, a statement about which the sequential construction carries no information.

Theorem 6 shows that, at the median scale, attainable overlap profiles must concentrate at the corners. This corner-seeking behavior results from the successive quadratic depletion of free adjacencies. After a proportion *s* of a candidate genome has already been fixed, a fresh independent input genome supplies asymptotically only a proportion (1 −*s*)^2^ of adjacencies whose two extremities remain available. By contrast, a proportion 1 − *s* of the candidate genome remains to be filled. For 0 *< s <* 1, (1 −*s*) − (1 −*s*)^2^ = *s*(1 −*s*) *>* 0. Thus, once a substantial contribution has already been taken from earlier inputs, the supply available from each fresh independent input falls short of the remaining genome capacity by a strictly positive macroscopic amount.

The stochastic depletion of compatible choices is the mechanism behind the Haghighi–Sankoff corner-seeking dynamics. The free-edge law exhibits this depletion at first order, while the attainable-region and Pareto results describe the resulting geometry and the cost of compromise. Theorem 5 establishes the existence of mixed profiles, including noncorner boundary profiles, with high probability. Theorem 6 identifies the rigid median-scale geometry of this trade-off, since the only attainable profiles with total overlap one are the corners. These results give a rigorous signed-multichromosomal formulation of the corner-seeking phenomenon through attainable profiles and explain the emergence of noncorner compromise configurations at larger total distance.

The same attainable region *P*_*k*_ will reappear in the next section as the feasible payoff region of the gene-sharing game. Its Pareto boundary quantifies how moving away from the corner-directed regime toward increasingly balanced compromise reduces total attainable overlap and leads to anti-median behavior.

## 4 Game-theoretic interpretation of anti-medians

In this section we study the midpoints of a set of genomes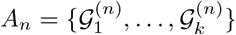, sampled independently and uniformly at random from G_*n*_. The midpoints of *G* _1_, …, *G* _*k*_ are those perfect matchings (genomes) which are equidistant from the input genomes *G* _1_, …, *G* _*k*_. In particular, within the sequential sampling framework of Section 3, we find the midpoints with the closest total distance to *A*_*n*_. These are called anti-medians and are obtained in an attempt to compromise the number of adjacencies shared from the input genomes [11]. Here, “anti-median” refers to the equal-compromise configuration on the Pareto boundary, not to a global maximizer of total breakpoint distance. We interpret this problem as a game of gene-sharing that is played by *k* players *G* _1_, …, *G* _*k*_. This reduces to an optimization problem that we solve and deduce that compromising results in an anti-median behavior. This fully explains the simulation observation regarding the appearance of anti-medians after compromising the gene adjacencies in [11].

Let 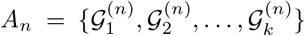 be *k* perfect matchings picked uniformly and independently at random from G_*n*_. We frame this as a game in which each genome _*i*_ plays the role of player *i*. The players *G* _1_, …, *G* _*k*_ cooperate.

Each player _*i*_, for *i* = 1, …, *k*, contributes a set *I*_*i*_ of its adjacencies, with |*I*_*i*_| ≈*α*_*i*_*n*, and the payoff of player *i* is *α*_*i*_. The contributions are made sequentially, and their order is essential to feasibility. For a permutation *σ* of [*k*], the players contribute in the order *G* _*σ*(1)_, …, *G* _*σ*(*k*)_. After the first *r* players have contributed, player *G* _*σ*(*r*+1)_ may contribute only adjacencies that are free with respect to those already selected. Thus the selected adjacencies form a (partial) matching that can be extended to a genome (a perfect matching).

For a fixed order *σ*, the asymptotically feasible payoff vectors are exactly the region 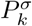 defined in (5). Allowing all possible orders therefore gives the feasible payoff region *P*_*k*_ defined in (6). For *α* = (*α*_1_, …, *α*_*k*_) ∈ *P*_*k*_, we define the total payoff by *u*(*α*) = *α*_1_ + ···+ *α*_*k*_.

The total payoff has a direct interpretation in terms of breakpoint distance.

For a constructed genome *G* whose normalized overlaps with the input genomes converge to *α*_1_, …, *α*_*k*_, as *n* → ∞, we have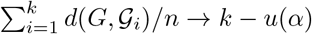. Thus maximizing the total payoff is asymptotically equivalent to minimizing the total breakpoint distance to the input genomes.

The players act as a team. They cooperate to choose a feasible profile and share the common payoff *u*(*α*), rather than pursuing their individual adjacency contributions. An unequal adjacency profile may therefore be optimal for the team if it maximizes the total payoff.

Since *u* is strictly increasing in every coordinate, every maximizer of the team payoff lies on the Pareto-optimal boundary of the feasible set. This boundary also contains the equal-compromise solution studied below.

### Definition 3.

*A feasible payoff vector α* = (*α*_1_, …, *α*_*k*_) *is Pareto optimal if no feasible vector* 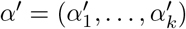*satisfies* 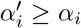 *for all i unless α*′ = *α*.

In other words, no player can gain more without making at least one player worse off. The set of all Pareto-optimal feasible vectors is called the Pareto optimal boundary of the game.

In the sequel, we find the Pareto optimal boundary *ϕ*_*k*_ of our gene-sharing game and show that the minimum payoff on *ϕ*_*k*_ occurs on its intersection with the diagonal { (*α*_1_, …, *α*_*k*_) : *α*_1_ = *α*_2_ = ···= *α*_*k*_ }, which is the compromiser of the payoff. We first study the case of two players (two random genomes) and then extend the results to general *k*.

### 4.1 Two players

For two players 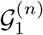 and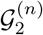, we may have two sampling mechanisms. Either player 1 first shares ⌊*α*_1_*n*⌋ of its adjacencies *I*_1_, and then player 2 can only share ⌊*α*_2_*n*⌋ of its free adjacencies with respect to *I*_1_, that enforces *α*_2_ ≤(1 − *α*_1_)^2^; or vice versa, player 2 shares first and player 1 shares next, enforcing *α*_1_ ≤(1 − *α*_2_)^2^. This means that the set of feasible payoff vectors is

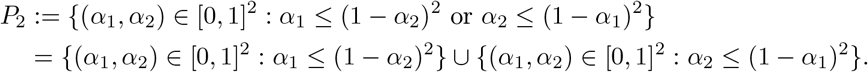

The first condition *α*_2_ ≤(1 − *α*_1_)^2^ is equivalent to either

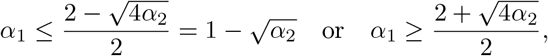

where the latter is impossible as it contradicts with *α*_1_ ∈ [0, 1]. This implies 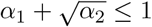. This and the same argument for *α*_1_ ≤(1 − *α*_2_)^2^ concludes *P*_2_ = {(*α*_1_, *α*_2_) ∈ [0, 1]^2^ :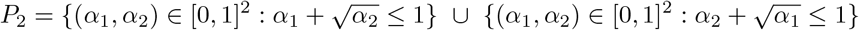:.

This is because the intersection of 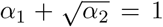 and 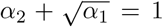 occurs at 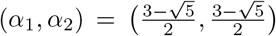 that comes as the intersection of both with the diagonal line *α*_1_ = *α*_2_, that is the solution to 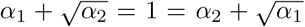(see sFigure 1). We have readily the following result.

**Figure 1:**
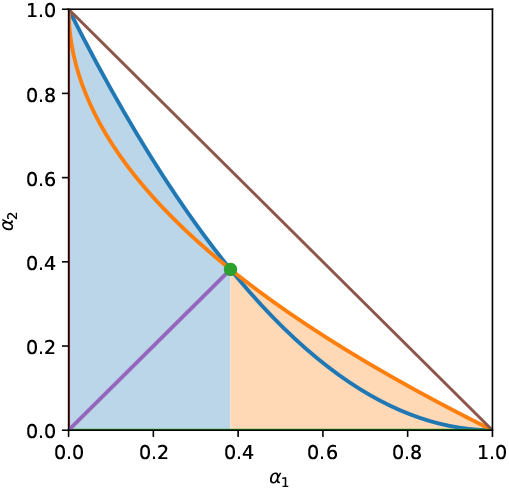
Feasible payoff region *P*_2_ (shaded) and Pareto boundary for the two-genome game. The boundary consists of the curves *α* = (1 *α* )^2^ and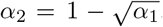. The diagonal segment *α*_1_ = *α*_2_ connects (0, 0) to their intersection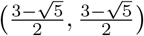, and the line *α*_1_ + *α*_2_ = 1 is shown for reference.

#### Theorem 7.

*For two-player games with players*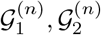, *the Pareto optimal boundary of feasible set of payoff vectors is the function ϕ*_2_ : [0, 1] → [0, 1],*defined by*

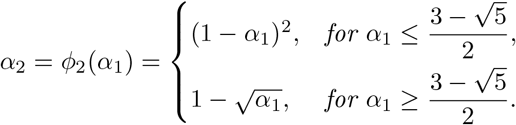

The set of feasible vectors *P*_2_ is the area surrounded by (inside and including) *α*_1_ = 0, *α*_2_ = 0, and *ϕ*_2_. As noted above, the optimal solutions of the gene-sharing game lie on *ϕ*_2_. Otherwise, at least one player can increase its payoff without making the other player worse off. So the first step for players *G*_1_ and *G* _2_ is to agree to choose their payoff vector on *ϕ*_2_. Therefore the optimization problem of finding the best and worst payoff *u*(*α*_1_, *α*_2_) = *α*_1_ + *α*_2_, reduces to:

#### Optimization Problem 1

(Median). *Find* (*α*_1_, *ϕ*_2_(*α*_1_)) *with α*_1_ ∈ [0, 1] *such that the function α*_1_ ↦ *α*_1_ + *ϕ*_2_(*α*_1_) *is maximized*.

#### Optimization Problem 2

(Anti-median). *Find* (*α*_1_, *ϕ*_2_(*α*_1_)) *with α*_1_ ∈ [0, 1] *such that the function α*_1_ ↦S*α*_1_ + *ϕ*_2_(*α*_1_) *is minimized*.

From the definition, it is clear that the function *α* _1_↦*α*_1_ + *ϕ* _2_ (*α* _1_ ) is strictly decreasing on 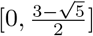 and strictly increasing on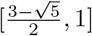. This implies that the solutions to the optimization problems (1) are (*α*_1_, *α*_2_) = (0, 1) and (*α*_1_, *α*_2_) = (1, 0). In other words the function *α*_1_ ↦*α*_1_ + *ϕ*_2_(*α*_1_), attains its maximum at *α*_1_ = 0 (i.e. (*α*_1_, *α*_2_) = (0, 1)) and *α*_1_ = 1 (i.e. (*α*_1_, *α*_2_) = (1, 0)) with maximum value *u*(0, 1) = *u*(1, 0) = 1 that corresponds to the approximate medians tending to the corners (input genomes 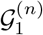 and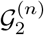).

On the other hand, the unique solution to the optimization problem (2) is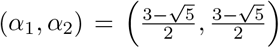. In other words, the function *α*_1_ ↦→ *α*_1_ + *ϕ*_2_(*α*_1_) attains its minimum at 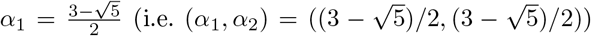 which corresponds to full compromise and anti-median.

Furthermore, as *α*_1_ ↦→ *α*_1_ + *ϕ*_2_(*α*_1_) is decreasing on 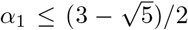 and increasing on 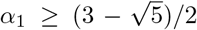, we can conclude that the more *G*_1_ and *G*_2_ compromise, the less the genome *G* constructed from their shared adjacencies behaves like a median, i.e. the more *G* behaves like an anti-median. This fully explains the observations from the simulation studies in [12, 11] for *k* = 2.

### 4.2 *k* players

We now extend the two-player analysis to an arbitrary number of players, using the feasible payoff region *P*_*k*_, defined in (6), and its Pareto-optimal boundary *ϕ*_*k*_ introduced above. We say that a permutation *σ* of [*k*] is a *witnessing order* for *α* if 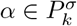. We first characterize Pareto optimality.

#### Theorem 8.

*Let α* = (*α*_1_, …, *α*_*k*_) ∈ *P*_*k*_. *Then α* ∈ *ϕ*_*k*_ *if and only if, for every i* = 1, …, *k*,

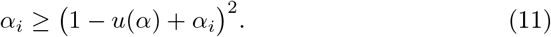

*Moreover, every α* ∈ *P*_*k*_ *satisfies* 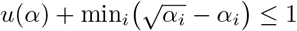, *and*

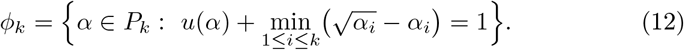

In view of (12), the normalized excess total breakpoint distance of a Pareto-optimal profile over the median value, 1 − *u*(*α*), equals 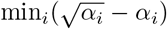.

We can now characterize the two extremal total-payoff configurations on the Pareto boundary. Let

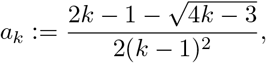

which is the smaller root of *a* = (1 − (*k* −)*a*) ^2^ and set *v*^*^ := (*a*, …, *a* ).

#### Theorem 9.

*For every k* ≥ 2, *the total payoff u on the Pareto-optimal boundary ϕ*_*k*_ *satisfies ka*_*k*_ ≤ *u*(*α*) ≤ 1. *The maximum value is* 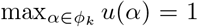, *and is attained precisely at the corners e*_1_, …, *e*_*k*_. *The minimum value is*

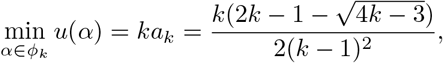

*and its unique minimizer is v*^*^ = (*a*_*k*_, …, *a*_*k*_). *Moreover*,

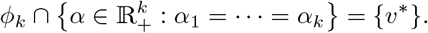

Theorem 9 identifies the two opposite configurations of the gene-sharing game. At the corners, the total payoff equals 1, corresponding to the corner-directed median behavior described in the previous section. At the other extreme, the unique symmetric Pareto point *v*^*^ distributes the payoff equally among all *k* players but has the smallest total payoff on *ϕ*_*k*_. Its corresponding total breakpoint distance to the *k* input genomes is asymptotically *kn*(1 −*a*_*k*_). Thus complete compromise comes at the greatest loss in total payoff along the Pareto-optimal boundary and gives the symmetric anti-median configuration. These two configurations form the endpoints of a broader trade-off between total payoff and balanced representation. We next examine how these compromise costs and attainable-profile geometry appear at finite genome sizes.

## 5 Numerical illustrations and finite-size behavior

The preceding sections characterize the region *P*_*k*_ of overlap profiles attainable by adapted sequential adjacency sampling and identify the corner and equal-compromise extremes of its Pareto-optimal boundary. In this section, we examine how these asymptotic results appear for genomes of finite size. We begin with the constrained-compromise problem for three input genomes and then consider equal compromise for a general number of inputs. We next illustrate the finite-size geometry of the overlap profiles generated by a broad collection of adapted sequential sampling policies. Finally, we compare the finite-size success probabilities of target profiles with the theoretical attainable region *P*_3_.

### 5.1 Finite-size cost of compromise

We first consider the constrained-compromise problem for three input genomes. For an attainable overlap profile *x* = (*x*_1_, *x*_2_, *x*_3_) ∈ *P*_3_, write 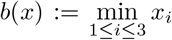, and, for *q* ≥ 0, define ℰ_3_(*q*) := max *u*(*x*) : *x* ∈ *P*_3_, *b*(*x*) ≥ *q* .

The interval on which this optimization problem is feasible is 0 ≤ *q* ≤ 1/4. Indeed, fix *x* ∈ *P*_3_ with *b*(*x*) ≥ *q*, and choose a permutation *σ* such that 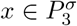. Set 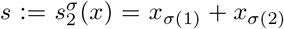. Since *b*(*x*) ≥ *q*, we have *s* ≥ 2*q*, while the final defining constraint of 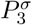 gives *q* ≤ *x*_*σ*(3)_ ≤ (1 − *s*)^2^. Consequently, 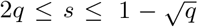, which is possible only when *q* ≤ 1/4. Moreover, *u*(*x*) = *s* + *x*_*σ*(3)_ ≤ *s* + (1 −*s*)^2^. The function *h*(*s*) := *s* + (1 −*s*)^2^ is convex, so its maximum on 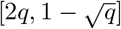 is attained at one of the two endpoints. Writing 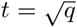, we have

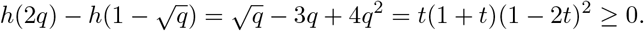

For 0 ≤ *q* ≤ 1/4, the profile *x*^*^(*q*) = (*q, q*, (1 −2*q*)^2^) belongs to 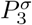 with *σ* the identity, since *q* ≤ (1 −*q*)^2^ and the final constraint holds with equality. Moreover, (1 −2*q*)^2^ −*q* = (1 −*q*)(1 −4*q*) ≥ 0, so *b*(*x*^*^(*q*)) = *q*.

The upper bound is therefore attained, giving ℰ_3_(*q*) = 2*q* + (1 −2*q*)^2^ = 1 − 2*q* + 4*q*^2^. The endpoints *x*^*^(0) = (0, 0, 1) and *x*^*^(1/4) = (1/4, 1/4, 1/4) correspond to a corner and equal compromise, respectively.

Since *µ*(*A*_*n*_)*/n* → 2 for three independent random genomes, a profile with total overlap *u*(*x*) has asymptotic normalized excess total breakpoint distance 1−*u*(*x*) above the median value. Therefore, the optimal constrained-compromise profile has

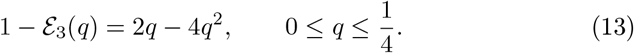

To examine this prediction at finite *n*, we used *n* = 100, 200, 400, 800. For these genome sizes, we generated 180, 140, 100, and 60 independent triples 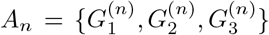, respectively. For each triple, we evaluated the construction over a regular grid of values *q* ∈ [0, 1/4]. For a given *q*, the adapted sequential construction, in the order 1, 2, 3, was assigned the integer target counts (⌊*nq*⌋, ⌊*nq*⌋, ⌊*n*(1 −2*q*)^2^⌋) . At each stage, the required number of adjacencies was sampled uniformly from those adjacencies of the currently revealed genome that were compatible with the partial matching already constructed. If fewer than the target number were available, all available adjacencies were retained and the construction continued. After the three stages, the unmatched vertices were paired uniformly at random to produce a complete genome, denoted by 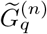. For every input triple we also computed the exact median value *µ*(*A*_*n*_). This can be done directly as a maximum-weight matching problem.

Figure 2 shows the empirical mean of 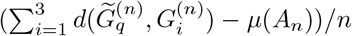 as a function of *q*, together with the asymptotic value (13). The finite-size curves closely follow the theoretical compromise-cost curve, and the discrepancy decreases as *n* increases. In particular, the two endpoints have the interpretations predicted by the theory. At *q* = 0, the optimal profile is a corner and the asymptotic excess vanishes, whereas at *q* = 1/4 the profile is the equal compromise (1/4, 1/4, 1/4) and the asymptotic normalized excess is 1/4. More generally, the figure shows at finite genome sizes the continuous increase in median cost produced by requiring larger minimum contributions from the three inputs.

**Figure 2:**
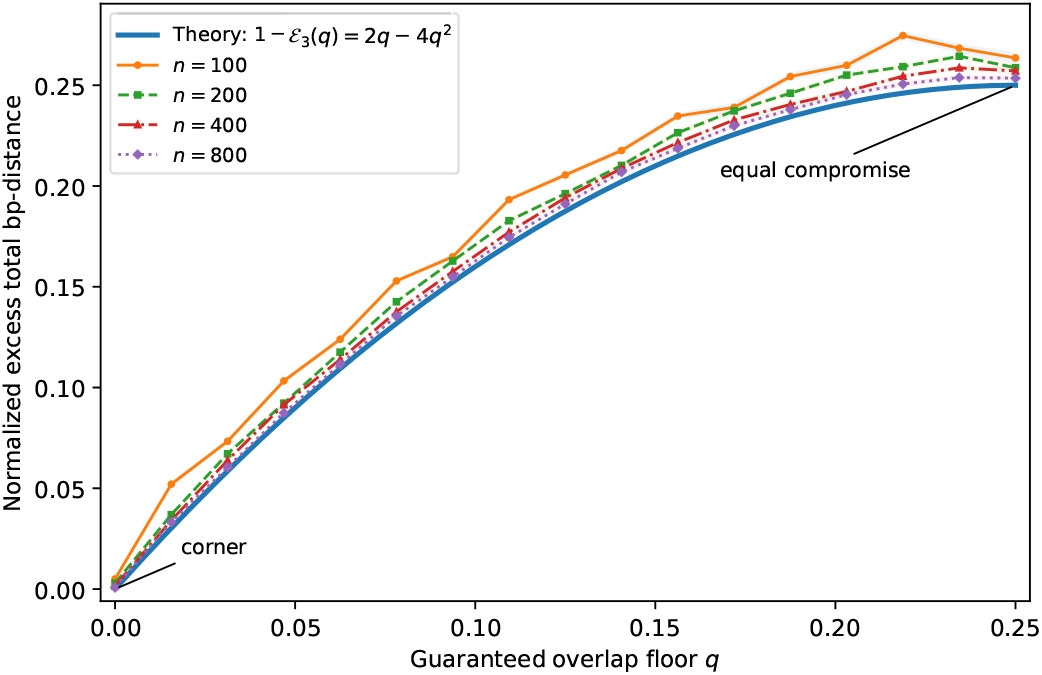
Finite-size cost of the optimal constrained compromise for *k* = 3. For *n* ∈ {100, 200, 400, 800}, the plotted curves show the empirical normalized excess as a function of the target overlap floor *q*. The solid theoretical curve is the asymptotic prediction 1 − ℰ_3_(*q*) = 2*q* − 4*q*^2^. The finite-size curves move toward the theoretical curve as *n* increases. The endpoints *q* = 0 and *q* = 1/4 correspond, respectively, to the corner solution and the equal compromise.

We next perform the analogous comparison for equal compromise with a general number of input genomes. Recall that the unique symmetric point minimizing *u* on the Pareto-optimal boundary of *P*_*k*_ is (*a*_*k*_, …, *a*_*k*_), where 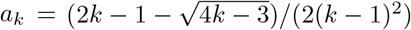. Its total payoff is *ka*_*k*_, so the asymptotic normalized excess over the median value is *ρ*_*k*_ := 1 − *ka*_*k*_.

For each *k* = 2, 3, 4, 5, 6 and *n* = 100, 200, 400, 800 we generated independent random input sets *A*_*n*_. We used 180, 140, 90, and 55 repetitions for *n* = 100, 200, 400, and 800, respectively. The target total number of selected input adjacencies was obtained by rounding *kna*_*k*_ to the nearest integer and distributing this number as evenly as possible among the *k* inputs. The genomes were revealed in the fixed order 1, …, *k*, and the same adapted construction as above was applied: the target number of currently compatible adjacencies was selected when available, all available adjacencies were retained in the event of a shortage, and the remaining vertices were paired at random. We denote the resulting complete genome by 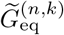.

Table 1 compares the empirical mean of 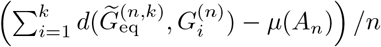 with the theoretical value *ρ*_*k*_. Across all values of *k*, the finite-size results show clear convergence toward the asymptotic prediction as *n* increases. By *n* = 800, the empirical excess is already quite close to 1 − *ka*_*k*_ throughout the range *k* = 2, …, 6, including the larger values of *k* for which finite-size effects are most noticeable. The agreement therefore provides a strong finitesize confirmation of the predicted cost of equal compromise and shows that the asymptotic description is already informative at moderate genome sizes. The general-*k* results show that this balanced-representation principle is not peculiar to three genomes: even as the number of inputs increases, the cost of requiring all lineages to contribute comparably remains quantitatively controlled.

**Table 1:** Finite-size cost of equal compromise. The theoretical column gives *ρ*_*k*_, the other entries are empirical means of the normalized excess total breakpoint distance. Standard errors were below 0.002 in all cases. There are 180, 140, 90 and 55 independent repetitions for *n* = 100, 200, 400, and 800, respectively.

| $k$ | $\rho_k$ | $n = 100$ | $n = 200$ | $n = 400$ | $n = 800$ |
| --- | --- | --- | --- | --- | --- |
| 2 | 0.23607 | 0.24350 | 0.24054 | 0.23981 | 0.23839 |
| 3 | 0.25000 | 0.26472 | 0.25729 | 0.25564 | 0.25400 |
| 4 | 0.24568 | 0.27172 | 0.25818 | 0.25408 | 0.25114 |
| 5 | 0.23799 | 0.27611 | 0.26046 | 0.25078 | 0.24323 |
| 6 | 0.22991 | 0.28750 | 0.25836 | 0.24653 | 0.23859 |

### 5.2 Finite-size overlap-profile geometry

The preceding calculations follow specific theoretically optimal compromise profiles. We next use a broader collection of adapted sequential policies to illustrate the range of overlap profiles that can arise at finite *n*. We take *k* = 3 and *n* = 400, and use the first independently generated input triple at this genome size as a representative instance.

For this experiment, a sampling policy consists of an ordering *σ* of the three input genomes together with a vector *γ* = (*γ*_1_, *γ*_2_, *γ*_3_) ∈ [0, 1]^3^ of stagewise sampling proportions, all fixed before the input genomes are used. We call the processing of *G*_*σ*(*r*)_ the *r*th stage of the sequential construction. At stage *r*, after *G*_*σ*(*r*)_ is revealed, suppose that *N*_*r*_ of its adjacencies are compatible with the partial matching selected in the preceding stages. The construction selects ⌊*γ*_*r*_*N*_*r*_⌋of these compatible adjacencies uniformly without replacement. After the three stages, the remaining unmatched extremities are paired randomly to complete the candidate genome. Thus, a policy specifies both the input sampling order and what proportion of the currently compatible adjacencies is selected at each stage, but it does not depend on the realized input genomes.

We considered all six input orders. For each order, we used a bank of 250 policies designed to cover a broad range of stagewise sampling behavior. Of these, 62 had their three sampling proportions drawn independently from Unif(0, 1), 62 from Beta(0.55, 0.55), and 61 from Beta(2, 0.8). A further 62 policies had their first two proportions drawn independently from Unif(0, 1) and the final-stage proportion fixed at one. Finally, for each order we included three deterministic reference policies: the fully saturated policy (1, 1, 1), the balanced reference policy (1/4, 4/9, 1), and the corner reference policy (0, 0, 1). This gives 250 policies per order and 1500 constructions in total. The uniform policies provide broad coverage of intermediate choices, the Beta(0.55, 0.55) policies place more weight near very small and very large sampling proportions, and the Beta(2, 0.8) policies favor relatively large fractions of the currently available adjacencies. Fixing the final-stage proportion at one provides additional coverage of policies that select all compatible adjacencies remaining at the last stage.

At stage *r*, after the corresponding genome has been revealed, suppose that *N*_*r*_ of its adjacencies are compatible with the partial matching constructed so far. If the policy specifies stage proportion *γ*_*r*_, we select ⌊*γ*_*r*_*N*_*r*_⌋of these adjacencies uniformly without replacement and then proceed to the next stage. After the three stages, the remaining unmatched extremities are paired at random to complete the candidate genome. Importantly, the profile plotted in Figure 3(a) is the *actual* overlap profile of the completed genome 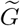,

**Figure 3:**
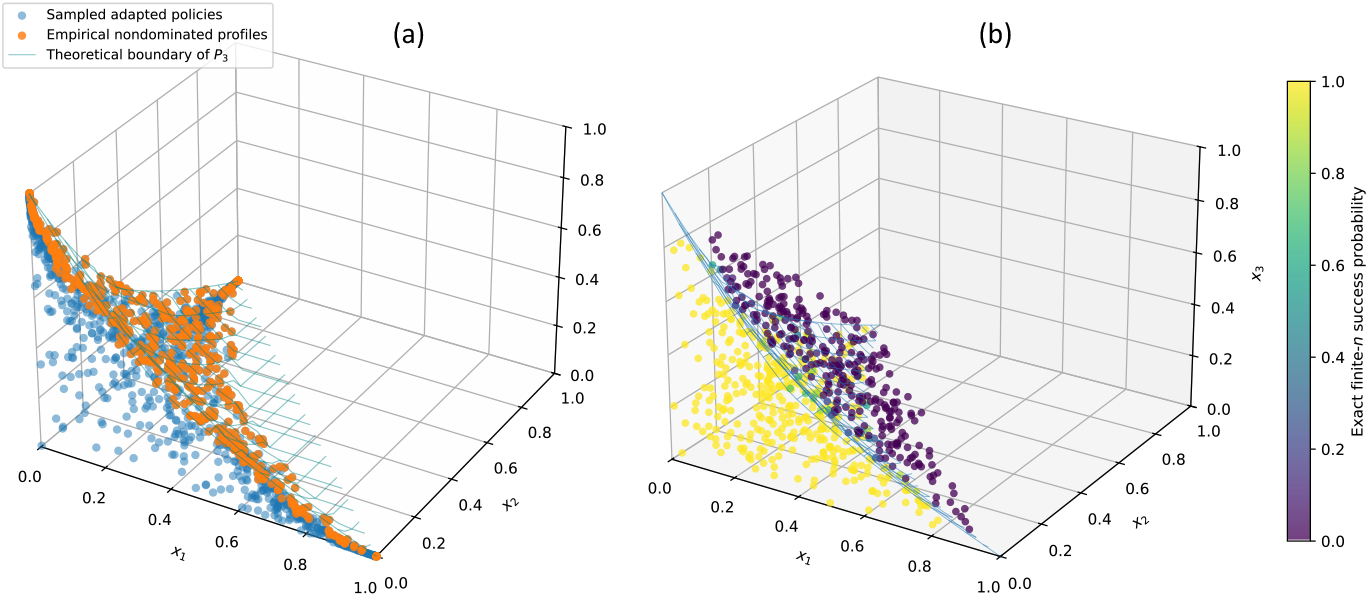
(a) Finite-size overlap profiles for a representative triple with *k* = 3 and *n* = 400. The light points are the actual profiles (14) obtained from 1500 adapted sequential constructions, with 250 policies for each of the six input orders. Highlighted points (in orange) are nondominated within this sampled collection. The wireframe is the theoretical upper boundary of *P*_3_. (b) Target-profile attainability for *k* = 3 and *n* = 400. The 700 target profiles are sampled uniformly from 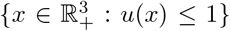 and are colored according to their exact finite-*n* success probabilities under the selected adapted order. The wireframe shows the theoretical surface separating *P*_3_ from its complement.

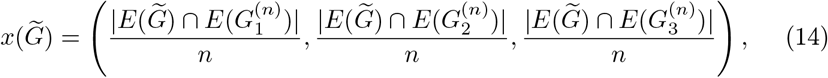

rather than the stage proportions used to generate it.

The lighter points in Figure 3(a) are the 1500 resulting profiles. The highlighted points are the empirically nondominated profiles: those for which no other profile in the sampled collection is at least as large in all three coordinates and strictly larger in at least one. The wireframe shows the theoretical upper boundary of *P*_3_. The sampled profiles reproduce the overall geometry predicted by the theory: the high-payoff extreme profiles occur near the corners, while more balanced profiles lie farther inward, and the empirically nondominated profiles trace the outer part of the attainable region. This gives a finite-size illustration of the trade-off between total payoff and compromise that underlies the preceding theoretical results.

The resulting cloud provides a finite-size view of the range and geometry of overlap profiles produced by a broad collection of adapted sequential sampling policies within the geometry predicted by *P*_3_. The different policy families populate both interior and more extreme parts of the feasible region, while the empirically nondominated profiles trace its outer portion particularly clearly. Because the policies were chosen to explore this geometry rather than to induce a particular probability distribution on profiles, the density of points reflects the breadth and design of the policy collection. The highlighted profiles identify the nondominated outcomes among the 1500 constructions considered here. Each profile records how the adjacency content of the constructed genome is distributed among the three inputs. Profiles near a corner are dominated by adjacency information from a single input genome, whereas profiles farther from the corners combine substantial contributions from several inputs. The cloud therefore gives a finite-size picture of the range of more input-dominated and more balanced reconstructions that can be obtained by sequentially combining adjacency information from the sampled genomes.

We conclude with a direct finite-size comparison with the attainable region *P*_3_. At *n* = 400, we uniformly sampled 700 target profiles from 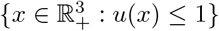. For each target *x*, we considered the six possible input orders and chose the order having the largest minimum margin in the defining inequalities of 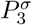. This choice is made from *x* alone and therefore does not use the subsequently revealed genomes. The given numbers of adjacencies were *m*_*i*_ = ⌊*nx*_*i*_⌋, *i* = 1, 2, 3.

For each target and chosen order, its finite-*n* probability of attaining all three given counts can be evaluated exactly. Conditional on having attained the preceding counts, the number of vertices remaining free is determined by those counts. The next genome is an independent uniform perfect matching, so the probability that sufficiently many of its adjacencies have both endpoints among the remaining free vertices is obtained by direct counting of perfect matchings. Applying this calculation successively over the three stages gives the exact success probability (4) shown in Figure 3(b).

The theoretical surface separates the region in which target profiles are asymptotically attainable by the adapted sequential construction with probability tending to one from the region in which the corresponding success probability tends to zero. Already at *n* = 400, the exact finite-size probabilities closely reflect this dichotomy: targets on the attainable side of the surface are predominantly assigned probabilities near one, whereas targets on the other side are predominantly assigned probabilities near zero. Thus the finite-size calculation provides a simple numerical illustration of the limiting region *P*_3_ without requiring an exhaustive search over sequential constructions.

## 6 Discussion

We have studied why breakpoint medians of independent random signed multichromosomal genomes, when constructed by sequential sampling, exhibit a strong tendency toward the input genomes rather than toward balanced intermediates. The probabilistic results show that, with high probability, the total breakpoint distance of a median is asymptotically close to its maximal scale (*k* −1)*n* and that a median can introduce only a vanishing proportion of adjacencies outside the input genomes. The attainable-profile analysis then identifies the first-order mechanism behind the corner-seeking behavior. More precisely, at the median scale, successive quadratic depletion prevents an adapted construction from retaining macroscopic contributions from multiple independent inputs. Almost all of its adjacencies must instead come from a single input, which may depend on the realization. This establishes a rigorous signed-multichromosomal attainable-profile formulation of the Haghighi–Sankoff phenomenon and gives a probabilistic explanation for the earlier observations in [9, 11].

The key mechanism is the progressive loss of compatibility among adjacencies. Once a proportion *s* of a candidate genome has been fixed, a newly revealed independent genome contains asymptotically only a proportion (1 −*s*)^2^ of adjacencies whose two extremities remain available. Thus the supply of usable adjacencies decreases faster than the unfilled fraction of the candidate genome. Repeating this loss across multiple inputs makes balanced overlap profiles increasingly difficult to attain. This shows that the tendency toward the corners is a consequence of the constraints on how adjacencies from different input genomes can be combined.

The game-theoretic formulation makes the resulting trade-off quantitative. Each input genome is viewed as a player whose payoff is the proportion of its adjacencies retained in the constructed genome. The total payoff is then directly related to total breakpoint distance. For arbitrary *k* genomes, the input-dominated corners maximize total payoff on the Pareto-optimal boundary, while the unique equal-compromise point minimizes it. For three genomes, the explicit constrained-compromise curve interpolates between these extremes and quantifies the loss in total payoff as the minimum contribution required from each input increases. In this way, medians, compromise solutions, and anti-medians arise as different choices within a common feasible-payoff geometry.

Our finite-size computations show that these asymptotic effects are already visible for genomes of moderate size. The observed compromise costs approach their theoretical limits as *n* increases, the sampled overlap profiles reproduce the predicted feasible geometry, and exact finite-size success probabilities closely distinguish target profiles lying inside and outside the asymptotically attainable region. The numerical results therefore provide finite-size support for the probabilistic and game-theoretic description and demonstrate that the predicted trade-off between minimizing total breakpoint distance and balancing contributions among the input genomes emerges clearly at moderate genome sizes.

These results also suggest a broader perspective on algorithm design. Different reconstruction goals naturally correspond to different regions of the feasible geometry. More precisely, minimizing total breakpoint distance favors median-type solutions, while objectives that emphasize balanced representation can be formulated through minimum-overlap constraints, maximin criteria, or target profiles along the Pareto boundary. The attainable-region analysis provides a quantitative framework for constructing and comparing such strategies, making explicit how different objectives navigate the trade-off between total similarity and balance among the input genomes.

The independent uniform model studied here provides a natural description of a deep-evolution regime, in which sufficiently many rearrangements have accumulated that genes, or larger conserved blocks used as the units of comparison, are effectively well mixed. In this regime, the random-genome model gives a useful asymptotic benchmark for understanding the geometry of breakpoint medians and the constraints on combining adjacency information from different inputs. From this perspective, our results describe the behavior that emerges after extensive evolutionary rearrangement. At shorter evolutionary distances, genomes descending from a common ancestor retain more shared adjacency structure, and the corresponding feasible geometry may interpolate between a closely related ancestral regime and the well-mixed regime analyzed here.

Overall, the results show that breakpoint optimality and balanced representation are distinct objectives whose tension is already encoded in the geometry of the genome space. The attainable-profile and game-theoretic viewpoints provide a common framework for quantifying this tension and for studying median, compromise, and anti-median reconstruction under alternative biological and computational objectives.

## Acknowledgments

Research partially supported by NSERC Discovery Grant to DS, RGPIN-2022-05212.

## A Proofs

### A.1 Proof of Theorem 1

*Proof*. For *G* ∈ G_*n*_, let 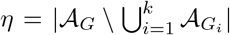. Since the sets 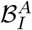, ∅ ≠ *I* ⊆ *A*, form a disjoint decomposition of 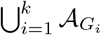, we have 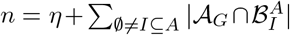. Moreover, every adjacency in 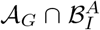 is shared by *G* with exactly |*I*| input genomes. Hence, for any *G* ∈ G_*n*_,

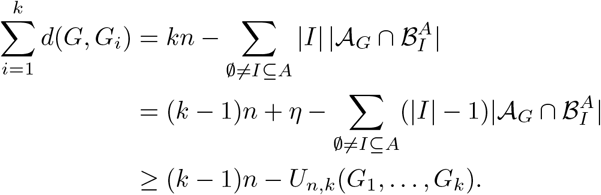

Taking the minimum over *G* gives the right-hand inequality in (1). For the other inequality, *µ*(*A*) ≤ min_*i*∈[*k*]_ ∑_*j*≠*i*_ *d*(*G*_*i*_, *G*_*j*_) = (*k* −1)*n* − *L*_*n,k*_(*G*_1_, …, *G*_*k*_), which gives the left-hand inequality.

### A.2 Proof of Lemma 1

*Proof*. Let *D*_*n*_ := {*G*_1_, …, *G*}_*k*_ are pairwise distinct . Since the genomes are independent and uniform in G_*n*_,

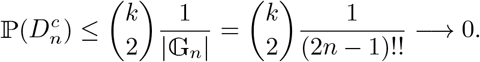

Fix distinct indices *i*_1_, …, *i*_*r*_ ∈ [*k*], with *r* ≥ 3, and let 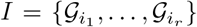. Conditional on 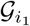, each of its *n* adjacencies belongs to all the remaining *r* − 1 independent random genomes with probability (2*n* − 1)^−(*r*−1)^. Therefore E|*A*_*I*_ | = *n/*(2*n* − 1)^*r*−1^ → 0. Since |*A*_*I*_ | is nonnegative and integer-valued, Markov’s inequality gives, as *n* → ∞, ℙ(|*A*_*I*_ | *>* 0) ≤ E|*A*_*I*_ | −→ 0. There are only finitely many choices of indices, since *k* is fixed. Together with 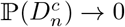, this proves the first assertion.

We next prove the first limit in (2). Fix *i* ∈ [*k*]. Conditional on *G*_*i*_, the random variables 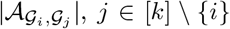, are mutually independent. Moreover, by invariance under relabelling, their conditional distributions do not depend on the particular value of *G*_*i*_. They are therefore mutually independent also unconditionally. By the pairwise Poisson limit recalled in Section 2, 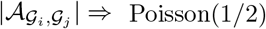 for *j* ≠ *i*. Since *k* is fixed,

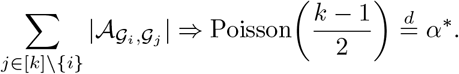

It remains to consider *U*_*n,k*_ = *U*_*n,k*_( *G*_1_, …, *G*_*k*_). Set 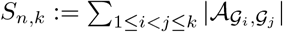. We first show that

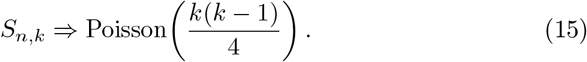

Put *N* = 2*n* and 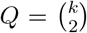. For an unordered pair *p* = {*i, j*} ⊂ [*k*], *i* ≠ *j*, and an arbitrary adjacency *ϵ* = {*a, b*} ⊂ ±[*n*], set 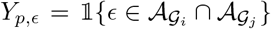. Then *S* = Σ_*p, ∈*_ *Y*_*p, ∈*_ . For *q* ≥ 1, let (*x*) = *x*(*x* − 1) · · · (*x* − *q* + 1) denote the falling factorial, and expand the factorial moment E[(*S*_*n,k*_)_*q*_] over ordered *q*-tuples of distinct pairs (*p, ϵ*). Consider first the terms for which the *q* selected edges are pairwise vertex-disjoint. If genome *i* is required to contain *d*_*i*_ of these edges, then Σ_*i*_ *d*_*i*_ = 2*q*, and a uniform perfect matching contains *d*_*i*_ prescribed disjoint edges with probability 1/((*N* − 1)(*N* − 3) · · · (*N* − (2*d*_*i*_ − 1))), where the empty product is 1. Thus the probability of such a term is *N* ^−2*q*^1 + *O*(*N* ^−1^)) . There are *Q*^*q*^ (*N* )_2*q*_/2^*q*^ such ordered choices. Their total contribution therefore converges to (*Q/*2)^*q*^ = (*k*(*k* − 1)/4)^*q*^ .

The remaining terms contribute *O*(*N* ^−1^). Indeed, if the *q* selected edges are distinct but not pairwise vertex-disjoint, there are *O*(*N* ^2*q*−1^) choices, and every compatible term has probability *O*(*N* ^−2*q*^). If instead only *h < q* distinct edges occur, there are *O*(*N* ^2*h*^) choices. Since the pairs (*p, ϵ*) in the factorial-moment expansion are distinct, some repeated edge must be associated with at least two distinct genome pairs and hence must occur in at least three genomes. Thus every compatible term imposes at least 2*h* + 1 distinct genome– edge requirements and has probability *O*(*N* ^−(2*h*+1)^). Hence, for every fixed *q*, E(*S*_*n,k*_)_*q*_ → (*k*(*k* − 1)/4)^*q*^ . By the method of factorial moments, (15) follows

On *D*_*n*_, an adjacency belonging to exactly *r* input genomes is counted 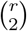 times in *S*_*n,k*_ and *r* − 1 times in *U*_*n,k*_. Consequently,

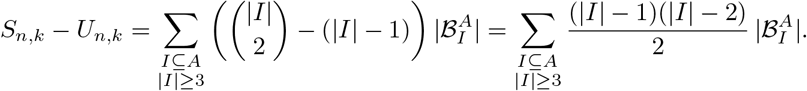

Since 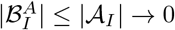 in probability, for every |*I*| ≥ 3, *k* is fixed, and 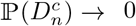, we have *S*_*n,k*_ − *U*_*n,k*_ → 0 in probability. Slutsky’s theorem [1] gives

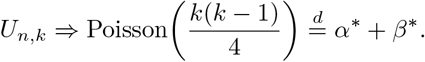

### A.3 Proof of Theorem 2

*Proof*. Set *X*_*n*_ = (*k* − 1)*n* − *µ*(*A*_*n*_) and let *D*_*n*_ be defined as in the proof of Lemma 1, where 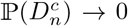. For any fixed *i* ∈ [*k*], on *D*_*n*_, Theorem 1 gives 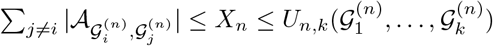. Hence, for every *r* ∈ ℤ_+_,

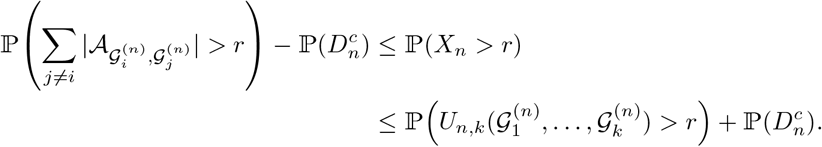

Thus, by Lemma 1,

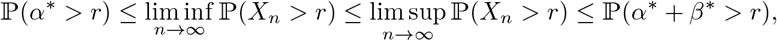

where we use convergence in distribution at the continuity point 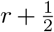, since all variables involved are integer-valued. Since on 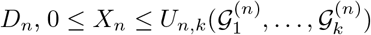, while 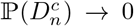, the tightness of 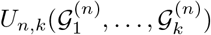 implies that (*X*_*n*_) is tight. Thus, by Helly’s selection theorem [1], every subsequence has a further subsequence converging in distribution. If 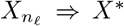, then *X*^*^ is also ℤ_+_-valued, and applying the preceding inequalities along this subsequence gives *α*^*^ ⪯_*st*_ *X*^*^ ⪯_*st*_ *α*^*^ + *β*^*^.

### A.4 Proof of Theorem 3

*Proof*. Let *X*_*n*_ := (*k* − 1)*n* − *µ*(*A*_*n*_) ≥ 0. By the previous theorem, the sequence (*X*_*n*_)_*n*≥1_ is tight. Hence, for every *ε >* 0, there exists *M <* ∞ such that30sup_*n*_ ≥ _1_ ℙ(*X*_*n*_ *> M* ) *< ε*. Let (*a*_*n*_)_*n*≥1_ be any diverging sequence of positive numbers with *a*_*n*_→∞ . Fix *δ* > 0. Choose *N* such that for all *n* ≥*N* we have *δa*_*n*_ ≥ *M* . Then, for *n* ≥*N*, ℙ(*X*_*n*_*/a*_*n*_ > *δ*) = ℙ(*X*_*n*_ > *δa*_*n*_) ≤ ℙ(*X*_*n*_ > *M* ) < *ε*. This completes the proof.

### A.5 Proof of Theorem 4

*Proof*. For *k* = 1, we have *τ*_*n*,1_(*G*_1_) = 0. For *k* = 2, the characterization of two-genome medians in Section 2 gives 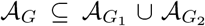 for every median *G*, and hence *τ*_*n*,2_(*G*_1_, *G*_2_) = 0. Thus, it remains to consider *k* ≥ 3. Let *A*_*n*_ = {*G*_1_, …, *G*_*k*_}, and let *D*_*n*_ be defined as in the pro f of Lemma 1, where 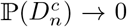. For every adjacency *e* on ± [*n*], let 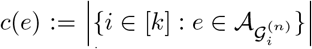. On *D*_*n*_ fix *G* ∈ *M* (*A*_*n*_) and set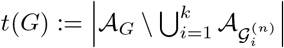. Then

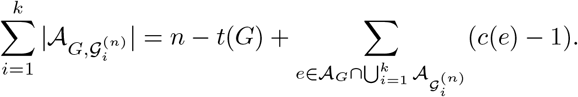

Since *G* is a median,

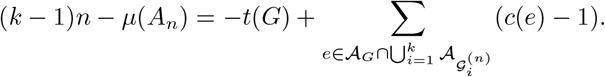

As (*k* − 1)*n* − *µ*(*A*_*n*_) ≥ 0, it follows that

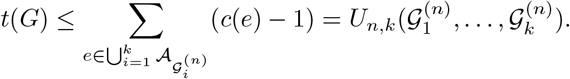

Since this bound holds for every *G* ∈ *M* (*A*_*n*_), on *D*_*n*_,

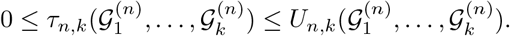

By Lemma 1, 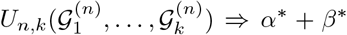, and hence this sequence is tight. Therefore, for every *ε, δ* > 0, there exists *M* < ∞ such that

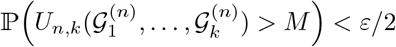

for all *n*, and, since *a*_*n*_ →∞, *δa*_*n*_ ≥ *M* for all sufficiently large *n*. Since also 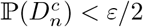 for all sufficiently large *n*, thus

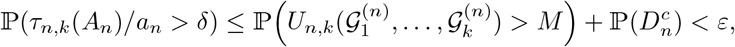

which proves the result.

### A.6 Proof of Proposition 1

*Proof*. Write *ℓ* = *n* − *m*, so that the complement of *R*_*n*_ has 2*ℓ* vertices. Since the uniform law on G_*n*_ is invariant under permutations of the vertices, the distribution of *F*_*n*_ depends on *R*_*n*_ only through its cardinality.

Each prescribed edge belongs to a uniform perfect matching with probability (2*n* − 1)^−1^, and each pair of disjoint prescribed edges belongs to it with probability ((2*n* − 1)(2*n* − 3))^−1^. Counting edges and ordered pairs of disjoint edges in the complement of *R*_*n*_ therefore gives, for *n* ≥ 2,

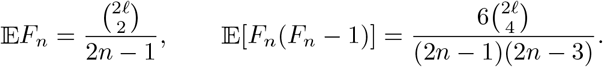

Here and below, binomial coefficients with upper argument smaller than the lower argument are understood to be zero. Consequently,

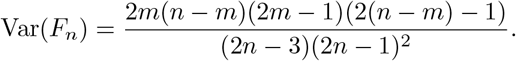

In particular, Var(*F*_*n*_) ≤*Cn* uniformly over 0 ≤*m*≤ *n*, for some absolute constant *C*. Moreover,

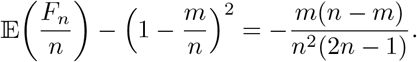

Thus, uniformly over 0 ≤*m* ≤*n*,

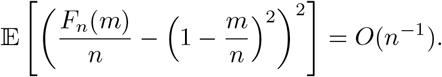

If *m*_*n*_*/n* → *x*, then (1 − *m*_*n*_*/n*)^2^ → (1 − *x*)^2^, and the result follows in *L*^2^.

#### Remark 3.

*One can obtain the asymptotics for the normalized number of edges G*^(*n*)^ *with both ends in R*_*n*_, *denoted by* 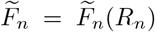 *or the normalized number of edges in with exactly one end in R*_*n*_ *and the other end in* ±[*n*] \ *R*_*n*_, *denoted by* 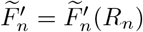. *For*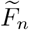, *we apply Proposition 1 to the set* ±[*n*]\*R*_*n*_, *instead of R*_*n*_, *and obtain*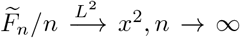. *For*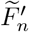, *we note that* 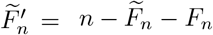, *hence*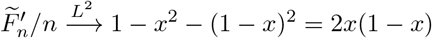 .

### A.7 Annealed equivalence of the two sampling laws

Fix *n*, and ***m*** = (*m*_1_, …, *m*_*k*_) with *M* := ∑_*i*_ *m*_*i*_ ≤*n*, and an order *σ*. Let *T*_***m***_ be the set of ordered tuples (*I*_1_, …, *I*_*k*_) of pairwise vertex-disjoint matchings on ±[*n*] with |*I*_*i*_| = *m*_*i*_. Write M(*N, a*) = (2*N* )!/((2*N* −2*a*)! 2^*a*^*a*!) for the number of *a*-edge matchings on 2*N* vertices, and put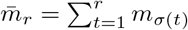, with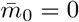.

Consider two experiments. In the sequential experiment, the independent uniform input genomes are revealed in the order *σ*, and at each stage the prescribed number of currently free edges is selected uniformly whenever possible. This is the prescribed-count scheme of Section 3 with uniform choices, and we write S for its success event and ***I*** = (*I*_1_, …, *I*_*k*_) for its selection, omitting the fixed index *n*. In the independent experiment, choose independently a uniform *m*_*i*_-edge subset *J*_*i*_ of each independent uniform input genome, and let C be the event that the blocks are pairwise vertex-disjoint. Then *ℒ* (***I*** | S) = *ℒ* (***J*** | C) = Unif(*T*_***m***_).

*Proof*. In the sequential experiment, conditional on success through stage *r* − 1, the occupied set has 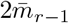 vertices and the next genome is independent and uniform. Invariance of the uniform law on G_*n*_ under the action of **S**_±*n*_ implies that the stagewise success probability *p*_*r*_ depends only on 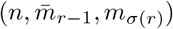 and that, conditional on stage success, the selected block is uniform among the 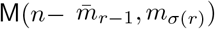 possibilities. Consequently, every ***t*** ∈ *T*_***m***_ has probability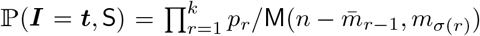, which is independent of ***t***. Conditioning on complete success gives the uniform law on *T* _***m***_. In particular, later success does not bias the earlier accepted blocks.

For the independent experiment, a uniform *m*_*i*_-edge subset of a uniform perfect matching is uniform among all *m*_*i*_-edge matchings on ± [*n*], by permutation invariance. Independence therefore gives every tuple of the prescribed sizes probability Π _*i*_ M(*n, m*_*i*_)^−1^. Conditioning on C gives the uniform law on*T*_***m***_.

Finally, successive counting yields

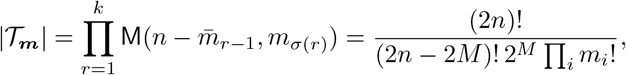

which also shows that the conditional law is sampling-order independent.

The equivalence is annealed and concerns only the selected tuple ***I***. It does not assert equality of the success probabilities or of the joint laws of the full input genomes and selections, and need not hold for arbitrary selection rules.

### A.8 Sequential sampling framework and proof of Theorem 5

We give the formal probabilistic formulation of Definition 1. Fix a permutation *σ* of [*k*]. For each *n*, let *ζ*_*n*,1_, …, *ζ*_*n,k*_ be auxiliary random variables, independent of the input genomes, that may be used to randomize the choices. Let 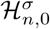 be the trivial *σ*-algebra and, for *r* = 1, …, *k*, let 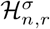 be the *σ*-algebra generated by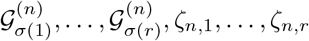 . Since the inputs are independent and the auxiliary variables are independent of them, 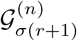 is independent of 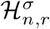 for every *r* = 0, …, *k* − 1.

#### Definition 4

(Formal version of Definition 1). *A* sequential sampling scheme *in the order σ is a stochastic process* 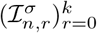 *with values in the finite set of*

*matchings on* ±[*n*], *adapted to*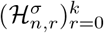, *with* 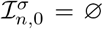*and* 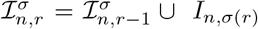, *where every edge of* 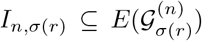 *is free with respect to the set* 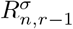 *of extremities covered by* 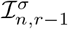.

In our constructions, 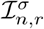 is obtained from 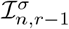by attaching a randomly chosen set *I*_*n,σ*(*r*)_ of free adjacencies of the current input, with a law depending only on 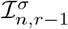 and the current input, so the process is Markov. The results here, however, require only adaptedness.

Each 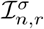is a matching, and adaptedness says exactly that the selection at stage *r* is a function of the inputs already considered and of the auxiliary randomness used so far. A prescribed-count scheme is the special case in which |*I*_*n,σ*(*r*)_ | = *m*_*n,σ*(*r*)_ at every stage until the first stage, if any, at which the current input has fewer than *m*_*n,σ*(*r*)_ free edges. At that stage the scheme fails, and *I*_*n,σ*(*t*)_ = ∅ for this and all later stages *t*.

Write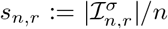, let 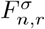be the number of edges of 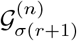free with respect to 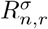, and set

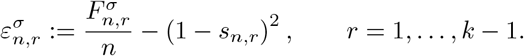

The set 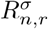is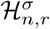 -measurable with 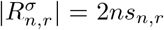 and 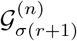is independent of 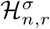. By the remark following Proposition 1, conditionally on 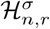the count 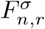 has the law of *F*_*n*_(*ns*_*n,r*_). Hence, for every *η >* 0, (3) gives

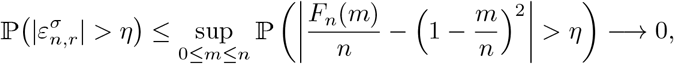

and, since *k* is fixed, a union bound yields

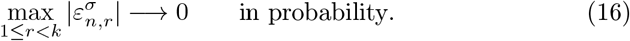

Since *I*_*n,σ*(*r*+1)_ consists of free edges and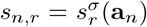, every scheme satisfies, without any success event,

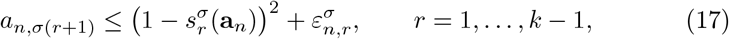

where **a**_*n*_ = (*a*_*n*,1_, …, *a*_*n,k*_). The passage to the limit uses the following deterministic lemma.

#### Lemma 2.

*For every ε >* 0 *there is δ >* 0 *such that every y* ∈ [0, 1]^*k*^ *with* 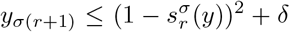 *for r* = 1, …, *k* − 1 *satisfies* dist 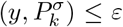 *and, if moreover u*(*y*) ≥ 1 − *δ, also* dist(*y*, {*e*_1_, …, *e*_*k*_}) ≤*ε*.

*Proof*. Suppose the first statement fails for some *ε >* 0. Then there are *y*^(*j*)^ ∈ [0, 1]^*k*^ satisfying the hypothesis with *δ* = 1*/j* and dist 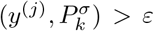. A sub-sequence converges to some *y* ∈ [0, 1]^*k*^, which by continuity satisfies the constraints with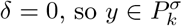. Since dist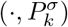 is 1-Lipschitz, this contradicts Dist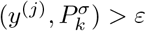.

The second statement follows similarly. A limit point *y* belongs to 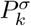 and satisfies *u*(*y*) ≥ 1, hence *u*(*y*) = 1 by Remark 2. Applying the quadratic constraints backward through the order *σ* forces *y* to be a corner.

*Proof of Theorem 5*. We first prove (7). Given *ε >* 0, let *δ* be as in Lemma 2. On the event 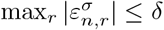 shows that **a**_*n*_ satisfies the hypothesis of the lemma, so dist 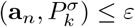. By (16), this event has probability tending to one. Necessity follows. Let *x* be *σ*-attainable, witnessed by target counts *m*_*n*_ := (*m*_*n*,1_, …, *m*_*n,k*_) and prescribed-count schemes with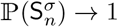. On 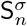we have **a**_*n*_ = *m*_*n*_*/n*. If dist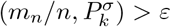, then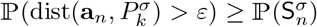, and by (7) this can happen only for finitely many *n*. Hence dist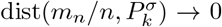, and since *m*_*n*_*/n* → *x* and 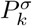 is closed,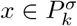.

We next prove sufficiency. Suppose first that all the inequalities defining 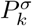 are strict at *x*, and choose *m*_*n,i*_ = ⌊ *nx*_*i*_⌋ . By Remark 2, the total number of prescribed edges is at most *n*. At the first stage the required number can always be selected.

At stage *r* +1, conditional on success through stage *r*, the previously selected edges occupy exactly 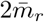extremities, and the next genome is an independent uniformly distributed perfect matching. Since 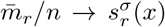and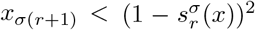, Proposition 1 implies that the conditional probability of having at least *m*_*n,σ*(*r*+1)_ free edges tends to one. Whenever this occurs, select the required number of free edges, using the auxiliary randomness if desired. Since *k* is fixed, applying this argument successively over the finitely many stages shows that the probability of success through all stages tends to one. Thus every profile satisfying the defining inequalities strictly is *σ*-attainable.

It remains to treat boundary points. Let 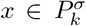 and choose a sequence *c*_*j*_ ↑ 1 with 0 < *c*_*j*_ < 1. Set *x*^(*j*)^ := *c*_*j*_*x*. Then *x*^(*j*)^ → *x*. If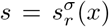, the defining constraint gives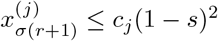, whereas

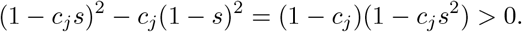

Hence all the defining inequalities are strict for *x*^(*j*)^, and the preceding argument gives a scheme with target counts 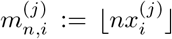whose success probability tends to one for each fixed *j*.

We may therefore choose integers *N*_1_ < *N*_2_ > ···, with *N*_*j*_ ≥ *j*, such that, for every *j* and every *n* ≥ *N*_*j*_, the scheme with target counts 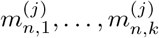 succeeds with probability at least 1 − 1*/j*. For *n* ≥ *N*_1_, define *j*(*n*) := max{*j* : *N*_*j*_≤*n}* and use the scheme with target counts 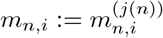. For the finitely many smaller values of *n*, choose any admissible target counts and scheme. Since *j*(*n*) → ∞, the success probability tends to one. Moreover, for every *i*,

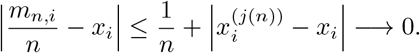

Thus *x* is *σ*-attainable. Taking the union over all permutations proves that the attainable region is exactly *P*_*k*_.

### A.9 Proof of Proposition 2

*Proof*. Let 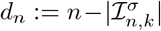be the number of edges added by the completion. Fix *i* ∈ [*k*] and put 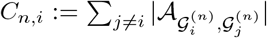, so that E*C*_*n,I*_ = (*k* − 1)*n/*(2*n* − 1) ≤ *k* − 1 by Section 2. Every edge of *I*_*n,i*_ belongs to both 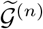 and 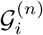, which gives the lower bound in (8). Any other common edge of 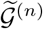 and 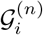 either lies in some *I*_*n,j*_ with *j* ≠ *i*, and hence in 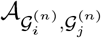, or is a completion edge. Writing *K*_*n,i*_ for the number of completion edges belonging to 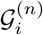, we obtain

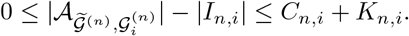

Conditionally on the inputs and all choices of the scheme, the completion is a uniform perfect matching on the 2*d*_*n*_ uncovered extremities. If *d*_*n*_ ≥ 1, each of the at most *d*_*n*_ edges of 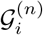 with both endpoints uncovered is included with probability (2*d*_*n*_ − 1)^−1^, so the conditional mean of *K*_*n,i*_ is at most *d*_*n*_/(2*d*_*n*_ − 1) ≤ 1. This is trivial when *d*_*n*_ = 0. Hence E(*C*_*n,i*_ + *K*_*n,i*_) ≤ *k*, and Markov’s inequality gives (8).

For the last assertion, let 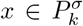 and take, by Theorem 5, target counts with *m*_*n,i*_*/n* → *x*_*i*_ and prescribed-count schemes with 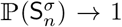. On 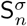 we have |*I*_*n,i*_| = *m*_*n,i*_, so |*I*_*n,i*_|*/n* → *x*_*i*_ in probability, and (8) gives the same limit for 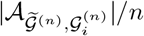. □

## A.10 Proof of Theorem 6

*Proof*. Let *x* ∈ *P*_*k*_ ∩ Δ_*k*_, and choose a permutation *σ* such that 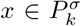. The final quadratic constraint gives 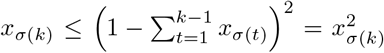, where the equality follows from ∑_*i*_ *x*_*i*_ = 1. Since 0 ≤ *x*_*σ*(*k*)_ ≤ 1, this forces *x*_*σ*(*k*)_ ∈ {0, 1}. If it equals one, *x* is a corner. Otherwise it equals zero, the preceding coordinates still sum to one, and the same argument applies to *x*_*σ*(*k*−1)_. Repeating this argument shows that exactly one coordinate equals one and all the others are zero. Conversely, every corner belongs to *P*_*k*_ ∩ Δ_*k*_, since for *e*_*j*_ we may choose an order with *σ*(1) = *j*. This proves the geometric identity.

Now let 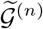 be as in the theorem, let *σ* be the order of the scheme, and write 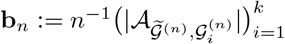. If 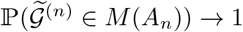, then (10) holds, since the excess is nonnegative and vanishes on 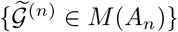 . Assume therefore (10). By (9),

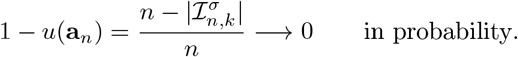

Given *ε >* 0, let *δ* be as in Lemma 2. By (17) and (16), with probability tending to one **a**_*n*_ satisfies both hypotheses of the lemma, and then dist(**a**_*n*_, {*e*_1_, …, *e*_*k*_}) ≤ *ε*. By Proposition 2, max_*i*_ |*b*_*n,i*_ − *a*_*n,i*_| = *O*_ℙ_(1*/n*), and the first convergence follows. Finally, 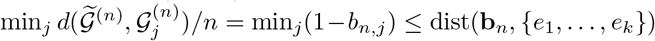. □

### A.11 Proof of Theorem 8

*Proof*. Suppose first that *α* ∈ *ϕ*_*k*_. Fix *i* ∈ [*k*], and let *σ* be a witnessing order for *α*. Delete player *i* from this order, while preserving the relative order of all remaining players. The resulting order witnesses the feasibility of the (*k* −1)-dimensional vector obtained from *α* by deleting *α*_*i*_: removing one of the preceding payoffs can only decrease the partial sums appearing in the sequential constraints and hence can only make the subsequent constraints easier to satisfy.

Suppose now that *α*_*i*_ *< (*1 − *u*(*α*) + *α*_*i*_)^2^. After the remaining *k* − 1 players have been taken in the order just described, append player *i* as the last player. The total payoff of the preceding players is ∑_*j*≠*i*_ *α*_*j*_ = *u*(*α*) −*α*_*i*_, so the final sequential constraint permits player *i* to receive any payoff up to

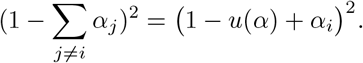

Hence *α*_*i*_ could be increased by a positive amount while all other payoffs remain unchanged. This would produce a feasible payoff vector strictly dominating *α*, contradicting *α* ∈ *ϕ*_*k*_. Therefore (11) holds.

Conversely, suppose that *α* ∈ *P*_*k*_ satisfies (11) for every *i*, and let *β* ∈ *P*_*k*_ satisfy *β*_*i*_ ≥ *α*_*i*_ for all *i*. Choose a witnessing order *σ* for *β* and put *j* = *σ*(*k*). Then

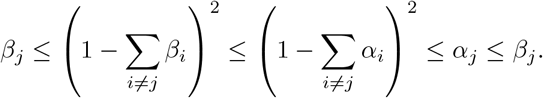

Since both sums are at most one and *s* ↦ (1 − *s*)^2^ is strictly decreasing on [0, 1], equality forces ∑_*i*≠*j*_ *β*_*i*_ = ∑_*i*≠*j*_ *α*_*i*_. Since also *β*_*j*_ = *α*_*j*_ and *β*_*i*_ ≥ *α*_*i*_ for every *i*, it follows that *β* = *α*, proving Pareto optimality.

For the scalar description, note that 1 − *u*(*α*) + *α*_*i*_ = 1 −∑_j≠*i*_ *α*_*j*_ ≥ 0 by Remark 2. Taking square roots, (11) is therefore equivalent to 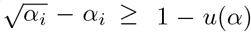, and so *α* ∈ *ϕ*_*k*_ if and only if 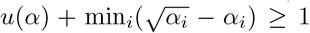. On the other hand, let *α* ∈ *P*_*k*_, let *σ* be a witnessing order and put *j* = *σ*(*k*). The last quadratic constraint gives 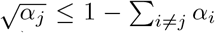, that is, 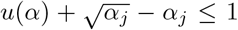. Hence 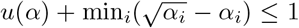 for every *α* ∈ *P*_*k*_, and equality characterizes *ϕ*_*k*_, which proves (12). □

### A.12 Proof of Theorem 9

*Proof*. By Remark 2, *u*(*α*) ≤ 1 for every *α* ∈ *P*_*k*_, and by the identity *P*_*k*_ ∩ Δ_*k*_ = {*e*_1_, …, *e*_*k*_} of Theorem 6, equality holds only at the corners. Every corner is Pareto optimal, since its total payoff is maximal, which proves the maximum assertion. Put 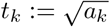, so that 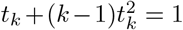. For *α* ∈ *ϕ*_*k*_, write *u* = *u*(*α*).

By (12) and the Cauchy–Schwarz inequality,

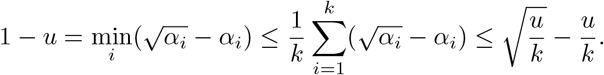

Setting 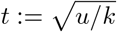 gives 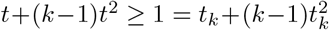. Since *t* ↦ *t*+(*k*−1)*t*^2^ is strictly increasing on [0, ∞), we have *t* ≥ *t*_*k*_, hence *u* ≥ *ka*_*k*_. If *u* = *ka*_*k*_, equalityholds in the Cauchy–Schwarz step, so all the *α*_*i*_ are equal to *u/k* = *a*_*k*_, that is, *α* = *v*^*^.

The vector *v*^*^ = (*a* _*k*_, …, *a* _*k*_ ) belongs to *P* _*k*_, since 1 − *ra* _*k*_ ≥ 1 − (*k* − 1)*a*_*k*_ = *t*_*k*_ *>* 0 and hence (1 − *ra*_*k*_)^2^ ≥ *a*_*k*_ for *r* = 1, …, *k* − 1. Moreover, 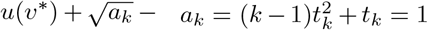, so (12) gives *v*^*^ ∈ *ϕ*_*k*_. Thus *v*^*^ is the unique minimizer.

Finally, if (*b*, …, *b*) ∈ *ϕ*_*k*_, then (12) gives 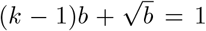. The left is strictly increasing in *b*, so *b* = *a*_*k*_, and *ϕ*_*k*_ meets the diagonal only at *v*^*^. □

## References

[1] Billingsley, P. Probability and Measure, 3rd ed. John Wiley & Sons, New York, 1995.

[2] Bryant, D. The complexity of the breakpoint median problem. Centre de recherches mathematiques (1998).

[3] Caprara, A. The reversal median problem. INFORMS Journal on Computing 15, 1 (2003), 93–113.

[4] da Silva, P. H., Jamshidpey, A., and Sankoff, D. Sampling gene adjacencies and geodesic points of random genomes. In RECOMB International Workshop on Comparative Genomics (2024), Springer, pp. 189–210.

[5] da Silva, P. H., Jamshidpey, A., and Sankoff, D. Identifying breakpoint median genomes: A branching algorithm approach. In 25th International Conference on Algorithms for Bioinformatics (WABI 2025) (2025), Schloss Dagstuhl–Leibniz-Zentrum für Informatik, pp. 18–1.

[6] da Silva, P. H., Jamshidpey, A., and Sankoff, D. Tree-based algorithms for breakpoint median and near-median genomes. preprint (arXiv) (2026).

[7] Feijão, P., and Meidanis, J. SCJ: a variant of breakpoint distance for which sorting, genome median and genome halving problems are easy. In International Workshop on Algorithms in Bioinformatics (2009), Springer, pp. 85–96.

[8] Fertin, G., Labarre, A., Rusu, I., Tannier, E., and Vialette, S. Combinatorics of genome rearrangements. The MIT Press, 2009.

[9] Haghighi, M., and Sankoff, D. Medians seek the corners, and other conjectures. BMC Bioinformatics 13, 19 (2012), S5.

[10] Jamshidpey, A., Jamshidpey, A., and Sankoff, D. Sets of medians in the non-geodesic pseudometric space of unsigned genomes with breakpoints. BMC Genomics 15, 6 (2014), S3.

[11] Larlee, C. A., Brandts, A., and Sankoff, D. Compromise or optimize? the breakpoint anti-median. BMC bioinformatics 17, Suppl 18 (2016), 473.

[12] Larlee, C. A., Zheng, C., and Sankoff, D. Near-medians that avoid the corners; a combinatorial probability approach. BMC Genomics 15, 6 (2014), S1.

[13] Sankoff, D., and Blanchette, M. The median problem for breakpoints in comparative genomics. Computing and Combinatorics (1997), 251–263.

[14] Sankoff, D., Sundaram, G., and Kececioglu, J. Steiner points in the space of genome rearrangements. International Journal of Foundations of Computer Science 7, 01 (1996), 1–9.

[15] Tannier, E., Zheng, C., and Sankoff, D. Multichromosomal median and halving problems under different genomic distances. BMC Bioinformatics 10, 1 (2009), 120.

[16] Xu, A. W. The median problems on linear multichromosomal genomes: Graph representation and fast exact solutions. Journal of Computational Biology 17, 9 (2010), 1195–1211.

[17] Xu, W., Alain, B., and Sankoff, D. Poisson adjacency distributions in genome comparison: multichromosomal, circular, signed and unsigned cases. Bioinformatics 24, 16 (2008), i146–i152.

[18] Zanetti, J. P. P., Biller, P., and Meidanis, J. Median approximations for genomes modeled as matrices. Bulletin of Mathematical Biology 78 (2016), 786–814.

